# Genomic connectivity and the spread of adaptive insecticide resistance alleles in *Anopheles arabiensis* from East Africa

**DOI:** 10.64898/2026.05.28.728377

**Authors:** Araya Eukubay, Kelly L. Bennett, Habte Tekie, Eric Lucas, Anastasia Hernandez-Koutoucheva, Fekadu Gemechu, Alistair Miles, Deriba Abera, Chris S. Clarkson, Lemu Golassa

## Abstract

Population connectivity and adaptive gene flow in disease vectors can shape the emergence and spread of insecticide resistance, with direct implications for control strategies such as insecticide spraying or the use of bed nets for malaria control. Using whole-genome sequencing, we first resolved the geographic population structure of the major but understudied malaria vector, *Anopheles arabiensis,* across the East African region, including the geographically diverse country of Ethiopia. We then assessed evidence for adaptive gene flow of insecticide resistance alleles across the region. Within Ethiopia, Central Rift Valley populations flanked by mountainous terrain were subject to restricted gene flow, although higher connectivity with the southwestern populations suggests an intermediate point of genetic exchange with the rest of Ethiopia. *Anopheles arabiensis* from western and northernmost Ethiopia were connected to populations from a similarly arid environment in Turkana - Kenya. Furthermore, broad-scale analysis revealed that populations from the rest of Kenya were connected with those from Uganda and Tanzania, but finer-scale analysis revealed more subtle structuring along the Rift Valley flanks, underscoring the role of landscape features in shaping patterns of gene flow. Adaptive gene flow analyses of diplotype clustering revealed that resistance alleles such as *Cyp6aa/p* and *Gste2* copy number variants (CNV) were widely spread across East Africa despite the geographical population structuring we observed. At the *Gste2* locus, *An. arabiensis* from Ethiopia and Kenya carried a newly annotated CNV spanning chromosome position 3R:28,596,832-28,606,222 linked to the non-synonymous SNP V47L, which was only at a low frequency in Kenya. Collectively, our findings demonstrate that *An. arabiensis* is subject to transboundary movement of resistance alleles, highlighting the need for coordinated cross-country vector management. Although areas of high connectivity may challenge genetic control technologies such as gene drives, more isolated populations may provide opportunities for targeted deployment.

**Significance:** Mosquitoes’ movement across landscapes determines how insecticide resistance spreads, yet the genomic connectivity of *Anopheles arabiensis* populations in East Africa is understudied. Using whole genome data, we show that geographic features such as the Rift Valley restrict gene flow in Ethiopia, with western and northernmost Ethiopia connecting to Turkana, while Kenyan populations outside Turkana connect further south with Tanzania and Uganda. Despite these geographic barriers, resistance variants at genes such as *Cyp6aa/p* and *Gste2* spread widely through adaptive gene flow, including a newly annotated CNV allele at *Gste2* in Ethiopia and Kenya. These findings demonstrate that resistance alleles move across national borders, underscoring the need for coordinated regional vector management. They also highlight that while broader connectivity may limit genetic control tools such as gene drive, more isolated populations offer opportunities for targeted deployment.

## Introduction

Mosquito disease vectors are globally distributed, with the pattern of their occurrence shaped by climate and natural or human-mediated dispersal (Abbasi 2025; Pabst et al. 2025). The geographic dispersal of mosquitoes has driven complex population structuring, which in turn directly impacts disease burden by influencing disease transmission dynamics across diverse ecological zones (Strauss et al. 2020). Connectivity among populations also influences how adaptive alleles, such as insecticide resistance variants, spread across landscapes (The Anopheles gambiae 1000 Genomes Consortium 2017; Clarkson et al. 2021; Odero et al. 2025). This movement of resistance alleles creates potential sources of spread and reintroduction after their suppression, which can be achieved using alternative insecticide treatments with a different selective pressure (Mwima et al. 2025). Conversely, population isolation can form localised resistance patterns that require tailored interventions (Mwinyi et al. 2025; Odero et al. 2025; Polo et al. 2025). Population connectivity also determines the feasibility of gene drive strategies, since their successful spread or containment depends on whether populations are highly connected or structured (Habtewold et al. 2025; Naidoo and Oliver 2025). Large-scale genomic studies of the major malaria vector *Anopheles gambiae* have shown extensive gene flow across West African populations, highlighting the risk of spread of adaptive variants, while also identifying distinct clusters in East Africa that may favour gene drive containment (The Anopheles gambiae 1000 Genomes Consortium 2017). In contrast, another major malaria vector, *An. arabiensis,* has remained comparatively underexplored, with genome-based population connectivity data sparse in Africa despite evidence of substantial heterogeneity in population structure and resistance variants in Kenya (Polo et al. 2025). Earlier studies using microsatellites, allozymes, and mitochondrial DNA haplotypes provided important evidence for weak population differentiation across large geographic distances (Lehmann et al. 1996; Donnelly et al. 2001; Nyanjom et al. 2003; Temu and Yan 2005; Muturi et al. 2010; Ng’habi et al. 2011; Maliti et al. 2014; Mustafa et al. 2021) but were based on few genetic markers with a low resolution to capture fine-scale dynamics. Modern whole-genome sequencing now enables simultaneous investigations of population connectivity, insecticide resistance variants, and evolutionary origins, offering unprecedented resolution to inform sustainable vector control strategies (The Anopheles gambiae 1000 Genomes Consortium 2017; Lucas et al. 2019; The Anopheles gambiae 1000 Genomes Consortium et al. 2020; Clarkson et al. 2021; Odero et al. 2025).

*Anopheles* mosquitoes in the *An. gambiae* species complex (*An. gambiae*, *An. Coluzzii* and *An*. *arabiensis)* and members of the *An. funestus* complex (s.l.) are the major malaria vectors across East Africa (Wiebe et al. 2017; Kweyamba et al. 2025). These species exhibit varied vectorial capacity and have adapted to diverse ecological conditions (Kahamba et al. 2022; Mwima et al. 2023). Unlike *An. gambiae* s.s., *An. coluzzii* and *An. funestus s.s*, which are primarily adapted to humid environments with abundant rainfall and permanent water bodies (Gimonneau et al. 2012; Msugupakulya et al. 2023), *An. arabiensis* extends into arid and semi-arid environments and is frequently associated with higher altitude habitats (Kirby and Lindsay 2004; Eba et al. 2021; Ashine et al. 2024; Esayas et al. 2024). Its ecological and behavioural plasticity, including exophagic and zoophilic tendencies, has enabled the species to expand its distribution across a wide geographical area in East Africa (Kreppel et al. 2020; Ashine et al. 2024), sustaining malaria transmission in regions where the other major vectors are less prevalent (Kitau et al. 2012; Lwetoijera et al. 2014; Killeen et al. 2016). In Ethiopia, *An. arabiensis* is recognised as the principal malaria vector (Eligo et al. 2024) and is found across the whole country, including the western and southeastern peripheral lowlands (Barasa et al., 2025; Chanyalew et al., 2022; Messenger et al., 2017; Woyessa & Yewhalaw, 2025), the Great Rift Valley (Gari et al. 2016; Eba et al. 2021) and the highland areas (Esayas et al. 2024). Its wide distribution reflects its ability to exploit diverse breeding habitats, including clear sunlit waters in permanent and temporary streams, hoofprints, artificial dams, roadside puddles, borrow pits, and rain pools (Getachew et al. 2020; Tarekegn et al. 2022). Although *An. arabiensis* is a major and widely distributed vector, studies of its population connectivity and genetic diversity in Ethiopia are lacking. Earlier microsatellite-based F_ST_ analyses detected low but statistically significant differentiation between populations from the northwestern highlands and the Central Great Rift Valley (Nyanjom et al. 2003), highlighting how Ethiopia’s diverse ecology may influence population structure. While population genetic data from other parts of East Africa provide some insight into regional *An. arabiensis* geographical population structure (Mwinyi et al. 2025; Polo et al. 2025), the extent of connectivity with the Ethiopian population is unknown. Addressing gaps in genomic data from across the region is critical for understanding vector population dynamics and guiding coordinated cross-country intervention strategies.

Resistance in *An. arabiensis* against insecticides used in indoor residual spraying (IRS) and insecticide-treated nets (ITNs) is widespread in Ethiopia (Alemayehu et al. 2017; Messenger et al. 2017). Insecticide resistance develops through two primary mechanisms: target-site resistance, where mutations alter the insecticide binding site (e.g kdr mutations in the voltage-gated sodium channel, *Vgsc*) (Martinez-Torres et al. 1998; Clarkson et al. 2021), and metabolic resistance, where overexpression or duplication of genes such as cytochrome P450s (Cyp450s), carboxylesterases (COEs) and glutathione S-transferases (GSTs) enables mosquitoes to break down insecticide before they act (Mitchell et al. 2014; Riveron et al. 2014; Lucas et al. 2024; Nagi et al. 2024). A genomic study of insecticide resistance-conferring genes across the country revealed spatial heterogeneity in the distribution of resistance markers: western Ethiopian populations are predominantly impacted by copy number variants (CNVs), associated with metabolic resistance, whereas northern populations show enrichment of target-site mutations (Eukubay et al. 2026). These findings highlight that resistance is not uniform across the country but shaped by local ecological and evolutionary pressures. However, it remains unknown how such resistance-conferring variants are shared across East Africa or influenced by barriers to population connectivity and gene flow. Understanding of these patterns is critical for anticipating and managing the spread of insecticide resistance, as the movement of resistance variants across borders may compromise the efficacy of insecticides to undermine malaria control.

In this study, we provide the first analysis of whole-genome sequence data of *An. arabiensis* within Ethiopia and in comparison to the broader East African region. We explore both population connectivity and the adaptive gene flow of insecticide resistance markers, thereby providing insights essential for tailored and cooperative vector control strategies across the region.

## Results

### Population connectivity and diversity of *Anopheles arabiensis* within Ethiopia

We constructed a neighbour-joining tree (NJT) based on genome-wide SNP data from *An. arabiensis* samples collected across multiple sites in Ethiopia to investigate population connectivity. The NJT revealed three distinct clusters: the first comprised western (Agnuak and Asossa) and northern (Werkamba) populations. The second included *An. arabiensis* from the Central Rift Valley, southern, northern (Raya-Azebo and Harbu), and southwestern populations. The third consisted of outlier samples from the southern site of Dilla (Supplementary Figure S2). Upon closer investigation, these outlier samples exhibited low genetic diversity and high homozygosity across a larger fraction of the genome compared to other samples from Dilla, possibly because they represent a group of related individuals with shared ancestry throughout the genome (Supplementary Figures S3). Consequently, we excluded them from further population structure analysis.

To investigate population connectivity further and corroborate findings from the NJT, we performed another form of dimensionality reduction via principal component analysis (PCA). Once again, populations from the western (Anguak and Asossa) and northernmost location (Werkamba) tended to group together and were partially separated from those in northern Ethiopia (Raya-Azebo and Harbu), the central Great Rift Valley, and the southern (Dilla and Arbaminch) locations, suggesting they are somewhat connected but experience restricted gene flow. In support, F_ST_ genetic differentiation was also lower between populations within the same NJT and PCA cluster (F_ST_ = 0.000 - 0.003) compared to populations in different clusters (F_ST_ = 0.005 - 0.008) (Supplementary Figure S4). Additionally, *An. arabiensis* populations from the southwestern site of Asendabo were found within both clusters, suggesting it experiences gene flow with geographically proximate populations in the southern region and Central Rift Valley as well as western Gambella (Figure 2). In support, F_ST_ genetic differentiation values were highest between Asendabo and the locations of Werkamba and Assosa (F_ST_ = 0.004), which are located furthest away in northern and westernmost Ethiopia, respectively. When the average PC1 scores were plotted by geographic location, we observed an east-west gradient in values correlated with longitude (Figure 3). We therefore used the genetic differentiation statistic F_ST_ calculated between population cohorts and geographic distances to assess isolation by distance (IBD) using the Mantel test. The findings show a significant relationship (Spearman r=0.43, p=0.013), consistent with IBD modulated by biological dispersal patterns. However, it is worth noting that *An. arabiensis* from Werkamba in the North and Asossa in the West have a similar genomic profile with an F_ST_ of 0.002 between the collection sites, despite being ∼622 km apart. Conversely, *An. arabiensis* from Werkamba are distinct from Raya-Azebo with an F_ST_ of 0.007, but are only 142 km apart within the same northern region.

**Figure 1.**
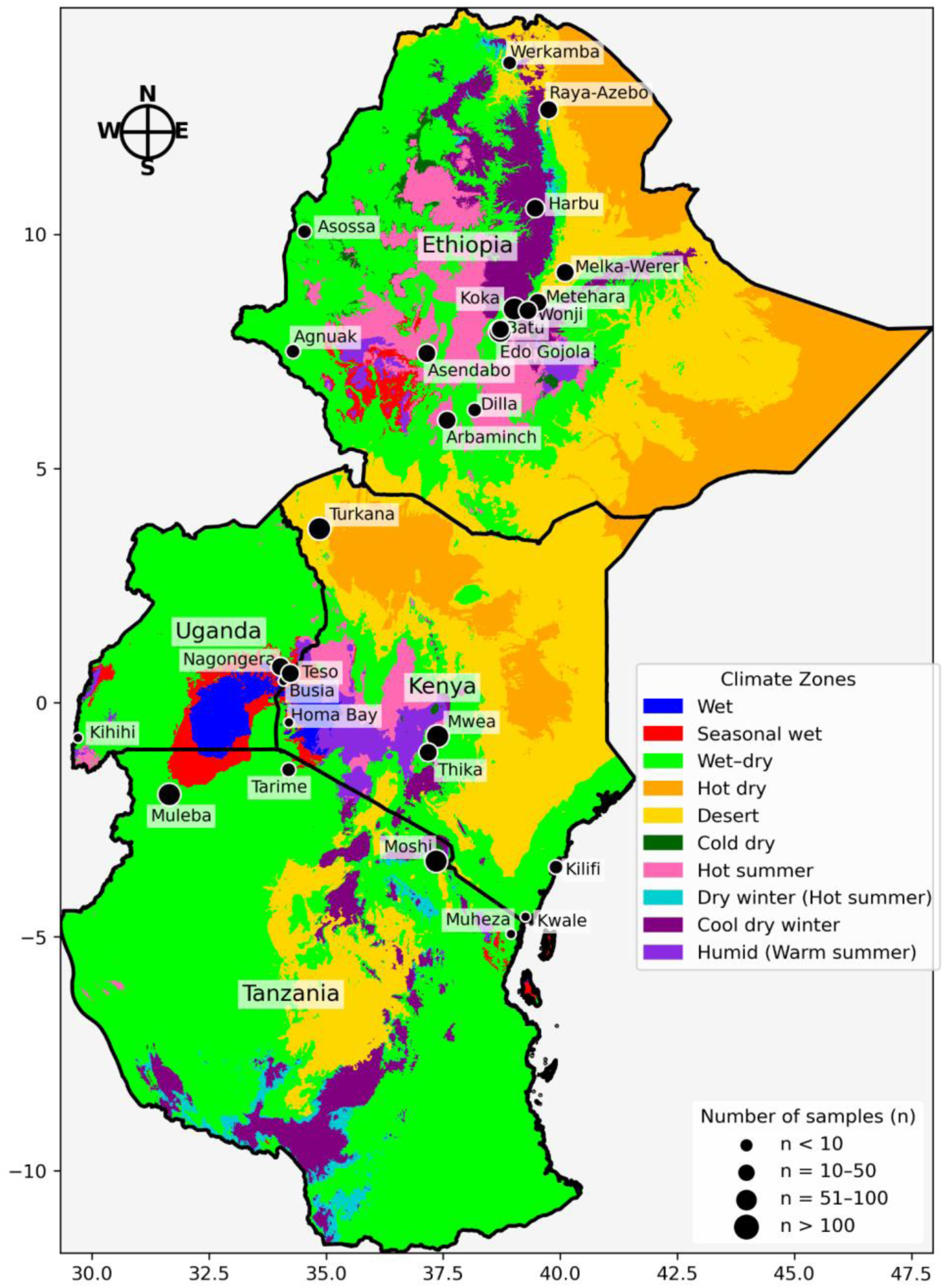
Map of sample populations in East Africa annotated with the Köppen–Geiger climate zones. The circle markers indicate sampling locations, with marker size scaled to the number of samples collected per site.

**Figure 2.**
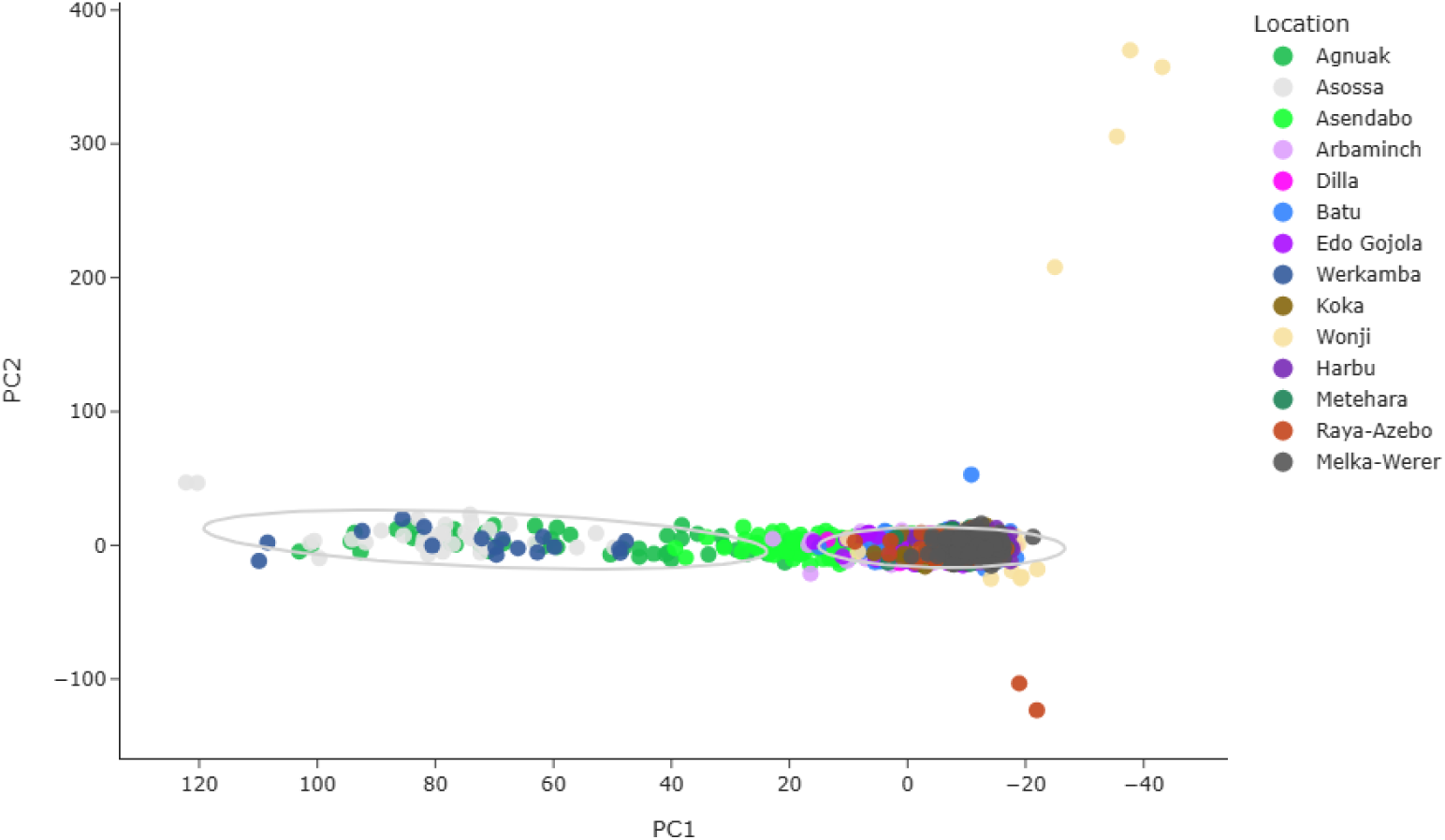
Principal component analysis (PCA) of *Anopheles arabiensis* in Ethiopia, coloured by sampling location. Samples are grouped into two clusters identified by K-mean clustering, with each cluster bounded by an ellipse derived from the covariance of its points to represent the 95% confidence interval.

**Figure 3.**
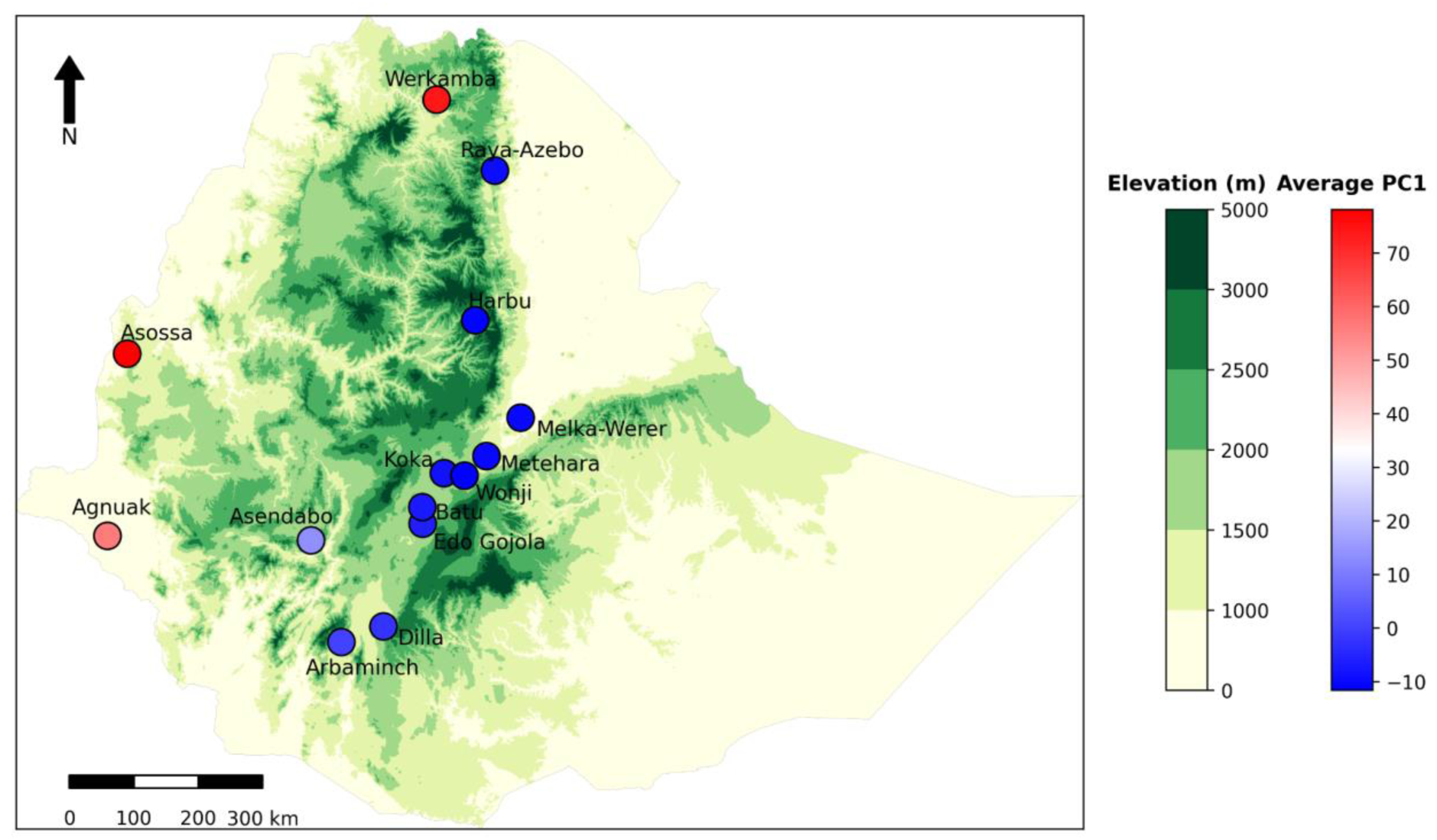
Mean PC1 scores of *Anopheles arabiensis* population in Ethiopia plotted on a geographic map overlaid with elevation. Sampling locations are represented by colour-coded markers, with a continuous blue-red gradient indicating relative genetic similarity. The right colour bar shows PC1 scale, while the left colour bar represents elevation ranges (altitude).

### Population connectivity of *Anopheles arabiensis* from East Africa

We extended our analyses beyond Ethiopia to include cohorts from Kenya, Tanzania, and Uganda using both NJT and PCA to evaluate broader transboundary population structure. The NJT analysis revealed that western and northern populations from Agnuak, Asossa, and Werkamba in Ethiopia are genetically similar to *An. arabiensis* from Turkana in northernmost Kenya, reflecting their confinement within comparable lowland Rift Valley ecological corridors (Supplementary Figure S5a). In contrast, Central Rift Valley and southern Ethiopian populations surrounded by the western and southeastern highlands also grouped independently, suggesting restricted gene flow with other East African *An. arabiensis* populations. All other populations from East Africa, including both inland and coastal regions, from northern Tanzania and southern central Kenya, formed a separate cluster on the NJT, suggesting they are more genetically similar to one another than to *An. arabiensis* from northern Kenya and Ethiopia.

Similarly, and in support of the NJT, the PCA revealed three clusters (Figure 4). Cluster 1 represented western and northern Ethiopia together with Turkana in northern Kenya, suggesting they are connected by gene flow. In support, F_ST_ analysis revealed western and northernmost Ethiopia (Agnuak, Asossa, and Werkamba) and Turkana - Kenya demonstrated minimal genetic differentiation (F_ST_=0.000-0.002). Cluster 2 included the Central Rift Valley and southern Ethiopian populations, which differed from the other Ethiopian and Turkana populations by an F_ST_ of 0.004 to 0.010. Finally, cluster 3 included all other East African populations, which differed from the other two clusters by an F_ST_ of 0.005 to 0.016 (Supplementary Figure S5b).

**Figure 4.**
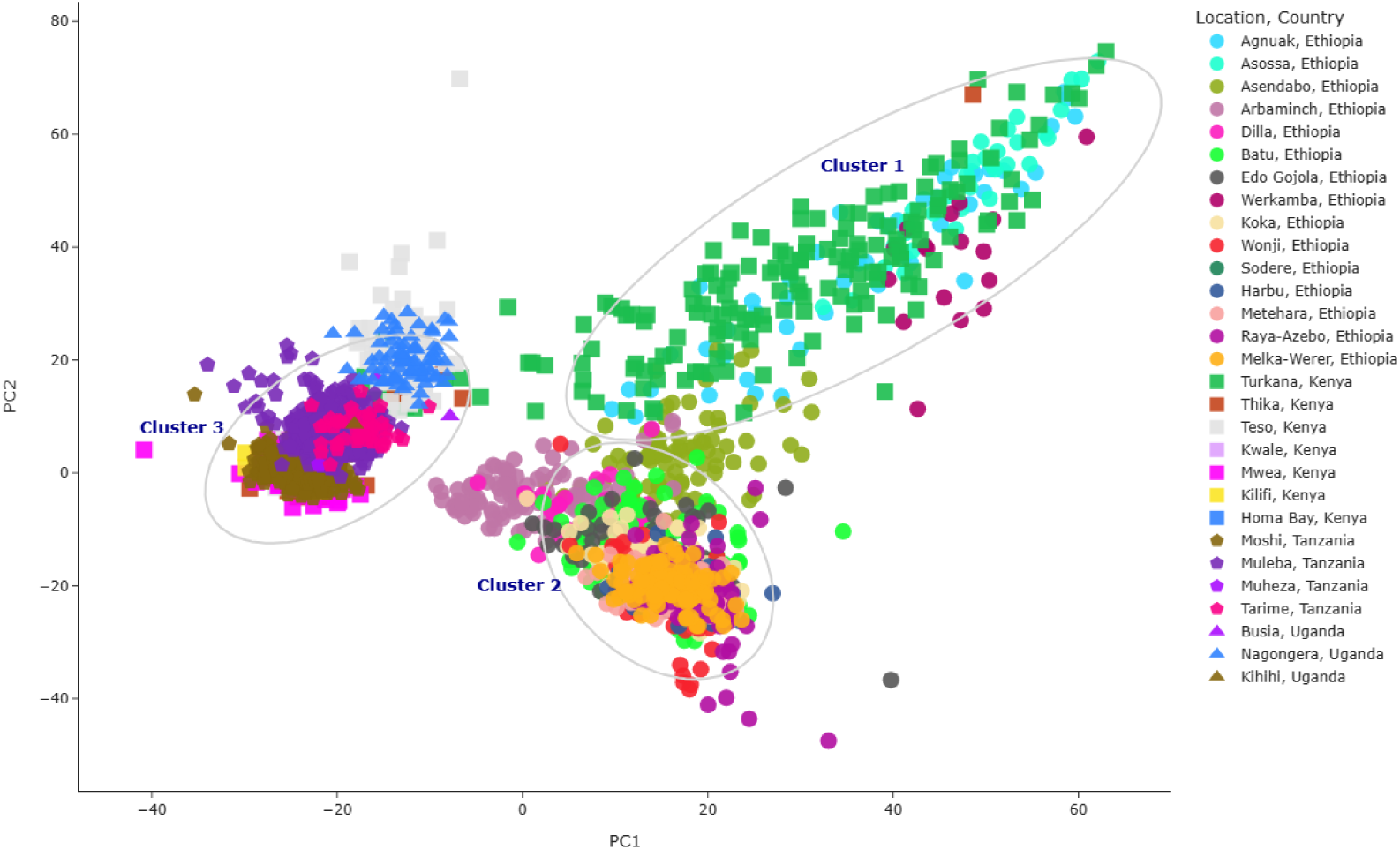
Principal component analysis (PCA) of *Anopheles arabiensis* in East Africa, coloured by sampling location. Samples are grouped into three clusters identified by K-means clustering. Each cluster is shown with an ellipse derived from the covariance of its points, scaled to represent the 95% confidence interval. **Cluster 1** corresponds to samples from western and northernmost Ethiopia and northern Kenya; **Cluster 2** represents samples from the Central Great Rift Valley and southern Ethiopia; **Cluster 3** encompasses all other East African samples, including those from Kenya, Tanzania, and Uganda.

To further resolve PCA cluster 3, we performed a separate analysis using only samples from Kenya (excluding Turkana), Tanzania, and Uganda. The PCA revealed three clusters, including Teso and Nagongera in western Kenya and Uganda (Cluster 1), Muleba and Tarime in northeastern Tanzania on the southern side of Lake Victoria (Cluster 2) and all other populations from central and coastal Kenya and Tanzania (Cluster 3) (Supplementary Figure S6). These clusters were supported by pairwise F_ST_ comparisons, with Teso—Nagongera separated from Muleba—Tarime by F_ST_=0.003-0.004. Teso and Nagongera were differentiated from central/coastal East Africa by a higher F_ST_ of 0.009-0.010, while Muleba and Tarime in northwestern Tanzania were also more differentiated from central/coastal East Africa with an F_ST_ of 0.003-0.004 (Supplementary Figure S5b). Overall, the PCA and F_ST_ values indicate finer-scale geographic structuring between populations located on the western versus the eastern side of the Rift Valley in East Africa as well as restricted gene flow between populations separated by Lake Victoria on the western side.

### Genomic diversity

To investigate whether populations of *An. arabiensis* across Ethiopia exhibit a similar demography, we calculated genetic diversity statistics for *An. arabiensis* sampled from different locations, years and for the minor and major malaria transmission seasons. The finding revealed uniformly high and stable nucleotide diversity (𝛑) and Watterson’s theta (𝛉_w_) across most cohorts with values ranging from 𝛑=0.0140 to 0.0149 and 𝛉_w_ = 0.0163 to 0.0187 (Figure 5). Tajima’s *D* was negative in all cohorts and more variable, ranging from -0.85 to -0.59. When values of Tajima’s D were lower, this was generally observed in the major transmission season, including for cohorts from the western locations of Agnuak and Asossa; the southern locations of Dilla and Arbaniminch; and northern Werkamba. The cohort for the 2023 major malaria transmission season in Dilla had noticeably higher values of genetic diversity (𝛑=0.0154 and 𝛉_w_=0.0198) and a more negative Tajima’s D (−0.940) than all other cohorts, suggestive of a larger or expanding population at this time. Furthermore, although the difference was slight, the western cohorts showed consistently higher diversity (𝛑=∼0.015 and 𝛉_w_=0.0190) and a lower Tajima’s D compared to most cohorts, potentially due to a larger population size or different demographic history. Conversely, values of Tajima’s D in the central Great Rift Valley were consistently higher, with values ranging from -0.59 to -0.67, suggesting a smaller population size or a slightly bottlenecked population.

**Figure 5.**
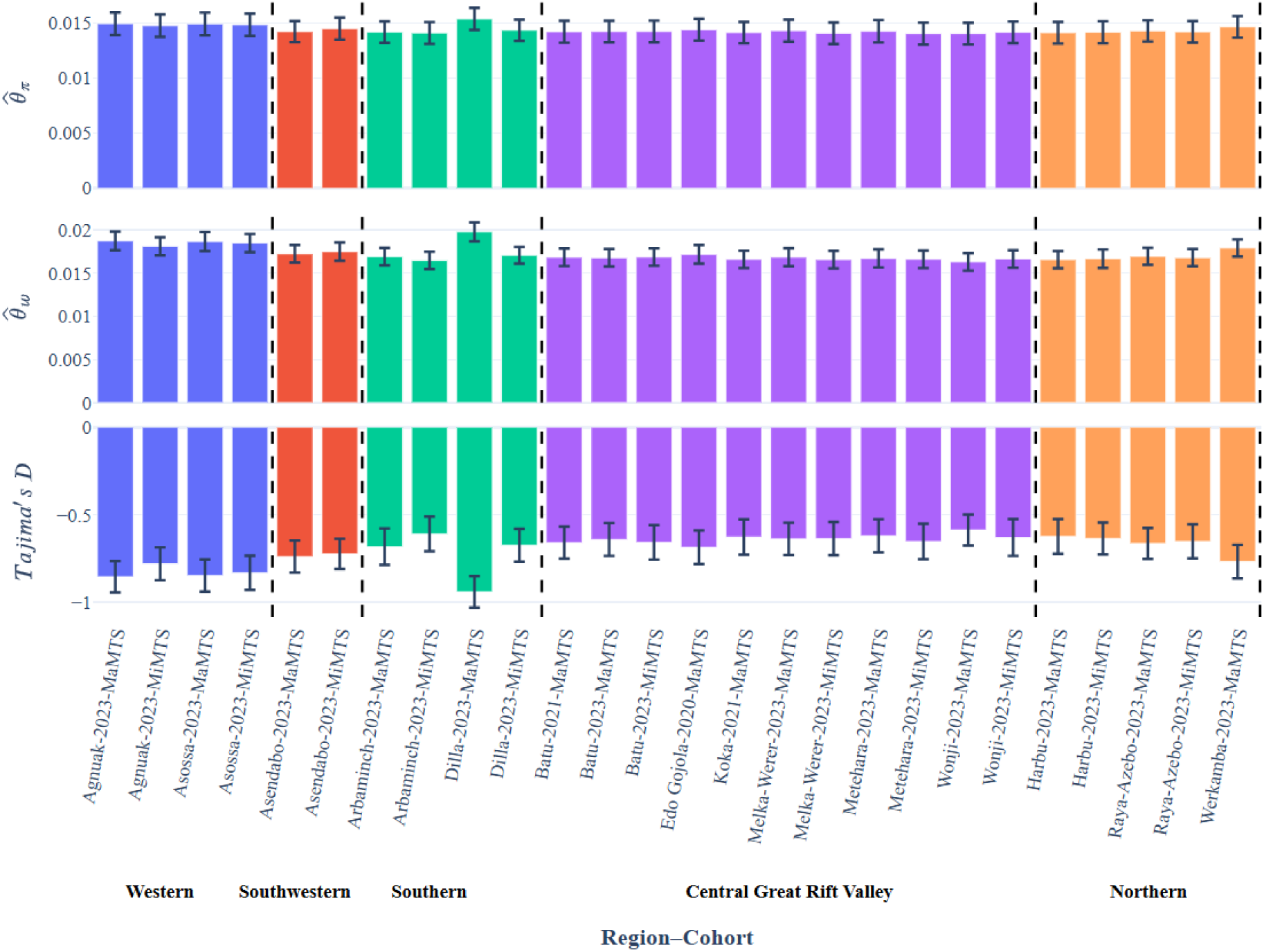
Genetic diversity of *Anopheles arabiensis* in Ethiopia. Nucleotide diversity (𝛉_𝛑_), Watterson’s estimator (𝛉_w_), and Tajima’s *D* by region-cohorts, year, and season. **MaMTS**: major malaria transmission season; **MiMTS**: minor malaria transmission season.

We also investigated the genetic diversity of *An. arabiensis* across East African countries, including Ethiopia, Kenya, Tanzania, and Uganda. The results showed a similar nucleotide diversity (𝛑) and Watterson’s estimator across most population cohorts, ranging from 𝛑=0.0135 and 𝛉_w_ = 0.0154 in Mwea, Kenya (2007), to 𝛑=0.0149 and 𝛉_w_ = 0.0187 in Agnuak, Ethiopia (2023). However, nucleotide diversity was much higher in Muleba in 2019, with values reaching 𝛑=0.0497 and 𝛉_w_=0.081 (Figure 6), while Tajima’s D was also low (−1.62), suggesting a comparatively large population compared to the previously sampled years. Tajima’s *D* was negative across all *An. arabiensis* cohorts sampled in East Africa, with the values ranging from -0.85 (Agnuak-2023, Turkana-2019) to -0.48 (Mwea-2007). Western Ethiopian populations (Agnuak and Asossa) and northern Kenya (Turkana) exhibited the highest diversity, whereas central Kenya (Mwea) consistently showed the lowest diversity estimator across multiple years. Similarly, central Ethiopian cohorts have higher Tajima’s *D* values, similar to cohorts from Mwea and Thika in Kenya. This highland region is characterised by intensive agriculture (Orondo et al. 2021) and strong evidence of metabolic resistance markers (Polo et al. 2025), suggesting that populations may be slightly bottlenecked due to insecticide exposure or reduced effective population size. Tajima’s *D* in the most recent Turkana cohort (2019) was comparable to western Ethiopian populations, indicating these populations may share similar demographic history.

**Figure 6.**
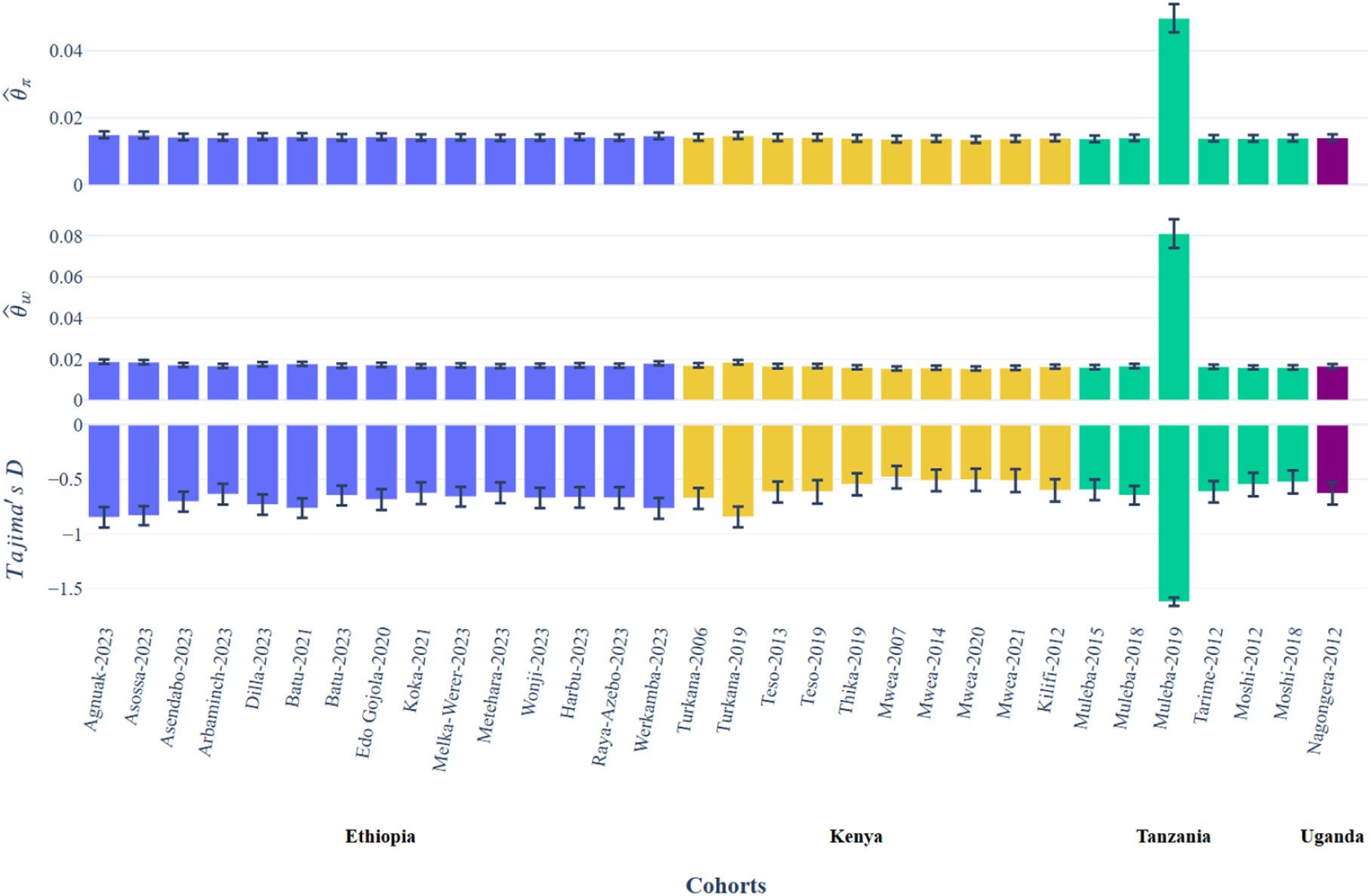
Genetic diversity of *Anopheles arabiensis* across East Africa. Nucleotide diversity (𝛑), Watterson’s estimator (𝛉_w_), and Tajima’s *D* are shown by location and year.

### Adaptive gene flow of molecular insecticide resistance

To investigate the adaptive gene flow of insecticide resistance variants, we constructed a hierarchical clustering dendrogram using diplotypes, i.e. multi-locus genotypes (Nagi et al. 2024) at known metabolic and target-site resistance loci previously identified as under selection in Ethiopia (Eukubay et al. 2026). Specifically, we investigated whether commonly found diplotypes with low heterozygosity at loci under selection were shared with *An. arabiensis* from other East African countries, including Kenya, Uganda, and Tanzania and assessed whether CNVs and non-synonymous SNPs were associated with the diplotype clusters. First, we applied diplotype clustering to the *Gste2* gene conferring metabolic resistance to organochlorines, organophosphates, and pyrethroids (Mitchell et al. 2014; Riveron et al. 2014). The result revealed that *An. arabiensis* from Ethiopia formed two diplotype groups with low heterozygosity at the *Gste2* gene. Cluster C1 included a small number of individuals from across Ethiopia characterised by both CNVs and a non-synonymous SNP unique to the cluster, V47L (Figure 7a). Cluster C2 with low heterozygosity was also unique to Ethiopia and included individuals with a high copy number but no non-synonymous SNPs associated with the cluster (Figure 7a). In addition, we observed another cluster of individuals from Ethiopia and only one sample from Kenya with a similar diplotype background and CNVs at *Gste2* (Cluster C3) but with elevated heterozygosity. In particular, the individuals within Cluster C3 were heterozygous for the SNP V47L and potentially represented individuals with both the C1 and C3 diplotype backgrounds we observed. To explore this notion further, we characterised the CNV architecture of the Ethiopian *An. arabiensis,* which were present in all three diplotype clusters. Two CNV alleles were identified, including the previously described Gstue_Dup12 and a novel duplication, Gstue_Dup17, spanning *Gste2*, *Gste1*, *Gste7* and *Gste3* (3R:28,596,832-28,606,222). Within the diplotype cluster C3, we found that 87% (167 out of 204) of Ethiopian individuals were heterozygous for both CNVs. Because CNV calls are only available at the genotype, not haplotype level, we applied Spearman’s correlation to identify phased SNPs that correlated with the presence of CNVs, which could be used as a proxy for the presence of the CNV at the haplotype level. Both CNV alleles showed a strong association with specific SNPs, including one at position 28,603,145 for Gstue_Dup12 (*ρ*t=0.998) and another at position 28,587,942 for Gstue_Dup17 (*ρ*=0.995). Notably, the SNP V47L at position 28,598,442 associated with diplotype cluster C1 was also highly correlated with Gsteu_Dup17 (*ρ*= 0.908). Plotting a haplotype dendrogram using these proxy SNPs further confirmed the presence of two major haplotype backgrounds in Ethiopian *An. arabiensis*, each linked to a distinct CNV allele (Supplementary Figure S7). Although V47L was consistently present in individuals with Gstue_Dup17, a minority of individuals without CNVs also carried the SNP on a divergent haplotype background, suggesting that V47L may have arisen prior to the evolution and spread of Gstue_Dup17 and that Gstue_Dup17 is more strongly associated with the low heterozygosity cluster than V47L.

**Figure 7.**
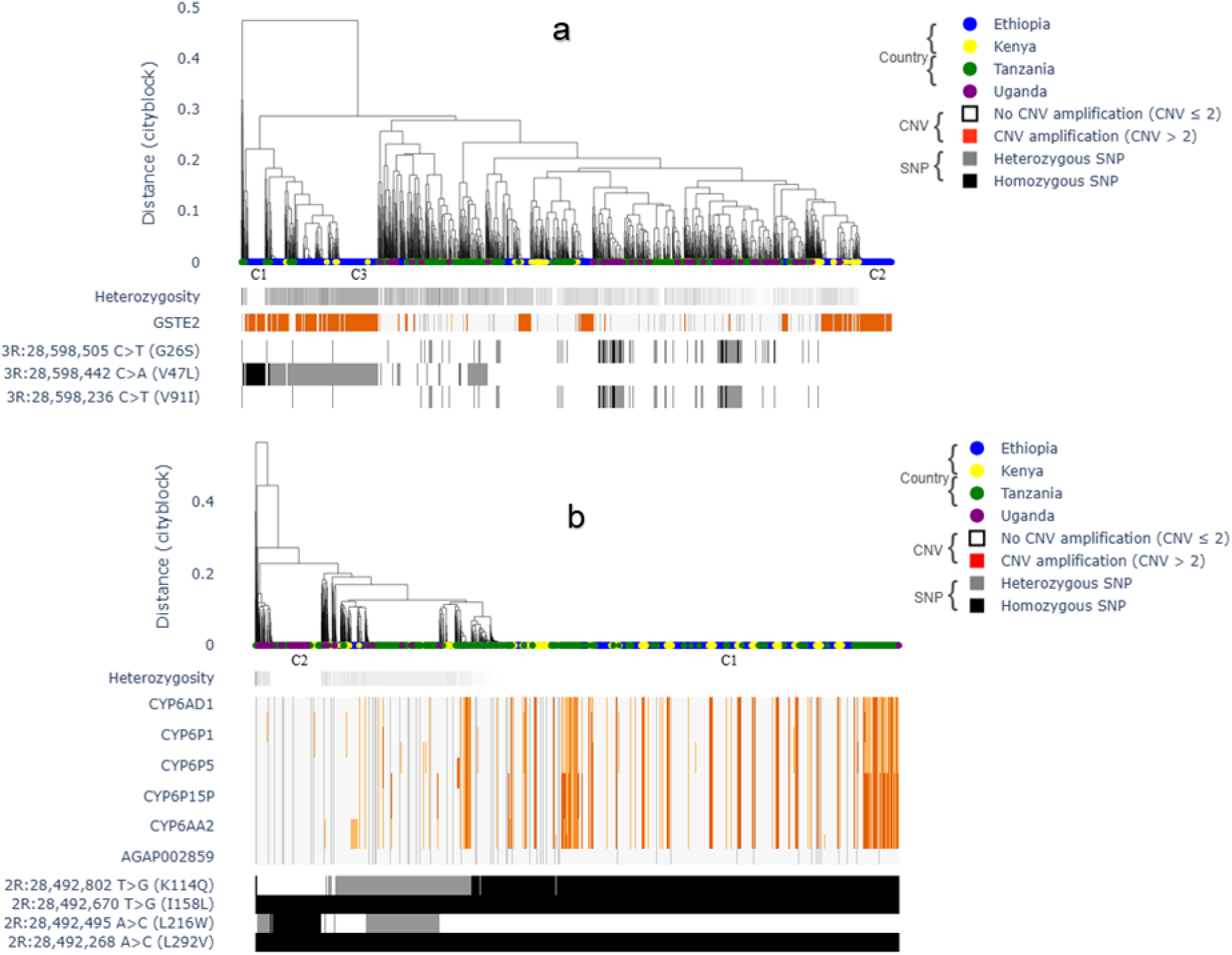
Diplotype clustering of East Africa *Anopheles arabiensis* at (**a**) *Gste2* and (**b**) the *Cyp6aa/p* gene cluster. Heterozygosity and copy number variation (CNV) are shown as colour bars below the dendrogram, with darker shades on the heterozygosity bar representing higher diversity and orange red color gradients on the copy number bar showing CNV amplification. Non-synonymous SNPs are plotted below the copy number bar, with gray to black color gradient, showing heterozygous and homozygous non-synonymous SNPs, respectively.

Next, we investigated the *Cyp6aa/p* gene cluster associated with pyrethroid resistance (Müller et al. 2008; Lucas et al. 2023; Nagi et al. 2024) and found a diplotype cluster (C1) with low heterozygosity, including many individuals with CNVs from all the East African *An. arabiensis* populations. We additionally observed a diplotype (C2) including *An. arabiensis* from Tanzania and Uganda only with a unique non-synonymous SNP (L216W) in *Cyp6p3*. This cluster exhibited marked reduced heterozygosity but no evidence of CNVs (Figure 7b), suggesting sample clustering could be driven by a point mutation rather than gene duplication. Because selective signals have also been reported in Ethiopia at the *Coejhe1-5e* gene cluster (Eukubay et al. 2026), and its presence is associated with insecticide resistance to organophosphates and pyrethroids in other East African countries (Abdalla et al. 2014a; Omoke et al. 2024), we investigated whether populations from these countries share a diplotype background with Ethiopian *An. arabiensis*. Although many individuals had a CNV spanning the *Coejhe3e* gene within the *Coejhe1-5e* cluster, similar to Ethiopian populations, we did not detect a cluster of individuals with low heterozygosity indicative of a swept haplotype (Supplementary Figure S8a).

We also applied diplotype clustering to the target-site resistance locus encoding the voltage-gated sodium channel gene (*Vgsc*). However, we did not observe a low heterozygosity cluster associated with the L995F allele, suggesting a particular diplotype background under selection (Supplementary Figure S8b). This may be because a selection signal has previously been observed only for a single cohort from northern Ethiopia (Eukubay et al. 2026), indicating selection could be geographically restricted. In addition, L995F appears to be relatively uncommon in East Africa, with most individuals observed as heterozygotes.

## Discussion

Historically, a number of studies investigating the population connectivity of *An. arabiensis* have revealed extensive gene flow and low genetic differentiation across large geographic areas, including across the high-altitude area of the Great Rift Valley in East Africa (Besansky et al. 1997; Kamau et al. 1999; Nyanjom et al. 2003; Hemming-Schroeder et al. 2020; Polo et al. 2025). Our analysis of population structure revealed unrestricted gene flow across much of Ethiopia and northern Kenya over a distance of approximately 1,209 km, with similar levels of observed genetic diversity indicative of a similar effective population size and shared demographic histories. However, we also found evidence for clear population structure in association with physical and/or ecological barriers to gene flow. In particular, we found reduced genetic diversity and lower connectivity of Ethiopian *An. arabiensis* in populations flanked by the Central Rift Valley, an area of high-altitude escarpments and mountainous terrains (Freilich et al. 2016; Behrends et al. 2024). As supported by a whole genome study within Kenya (Polo et al. 2025), we observed restricted gene flow between areas adjacent to high-altitude mountain ranges in Ethiopia and Turkana in northern Kenya and the rest of Kenya and Tanzania. We additionally observed more subtle geographic population structure between *An. arabiensis* located on either side of the Great Rift Valley and Lake Victoria in Kenya and Tanzania, which were previously reported to be important barriers to *An. arabiensis* gene flow (Polo et al. 2025). Importantly, it was noted that there are pronounced environmental differences between the regions impacted within Kenya which may also influence population connectivity (Polo et al. 2025). Additionally, we note that the areas of Ethiopia and Turkana we observed to be connected by gene flow occupy similar arid savannah environments with seasonal rainfall (Enyew and Hutjis 2015; Worku et al. 2022; Denje et al. 2023). Therefore, ecological similarities across the region may promote the dispersal of *An. arabiensis*. In contrast, populations in coastal Kenya, Tanzania, and Uganda experiencing more subtle geographic population structure are subject to more diverse ecological conditions, including humid subtropical coastal climates, temperate to semi-arid highlands, and irrigated agricultural systems (Luhunga and Songoro 2020; Zhou et al. 2022; Lawrence et al. 2023). In support of our findings, previous landscape genetic studies investigating population connectivity with microsatellite markers have shown that *An. arabiensis* dispersal is facilitated by cropland and aridity but restricted by ecological barriers, such as forests, lakes, and irrigated areas (Hemming-Schroeder et al. 2020). Interestingly, we observed that in Ethiopia, the southwestern populations appeared intermediate in genomic composition and F_ST_ values between the divergent groups in the Central Rift Valley and the rest of Ethiopia and northern Kenya. Findings raise the possibility that this region may function as a transitional zone where genetic exchange occurs. This pattern indicates the importance of ecological corridors in maintaining partial connectivity despite broader structuring (Bergey et al. 2020) and the need to comprehensively investigate the population connectivity of *An. arabiensis,* including across a fine geographical scale. The presence of intermediate populations is likely to facilitate the spread of adaptive alleles, including those conferring insecticide resistance (The Anopheles gambiae 1000 Genomes Consortium et al. 2020).

Our analysis of adaptive allele sharing revealed that key insecticide loci are shared across large geographical areas, consistent with the spread of adaptive variants under strong selective pressure (The Anopheles gambiae 1000 Genomes Consortium 2017; The Anopheles gambiae 1000 Genomes Consortium et al. 2020; Clarkson et al. 2021). For example, we observed the adaptive allele sharing of two major CNVs at the *Gste2* locus across Ethiopia and Kenya. The two CNVs had distinct diplotype and haplotype backgrounds and different tagging SNPs, suggesting they arose independently before their spread. The presence of Gstue_Dup17 as a homozygote in Ethiopia with the same diplotype background raises the possibility that this CNV may have originated there, with subsequent spread into Kenya, where the same duplication with a similar diplotype background was only found at a low frequency as a heterozygote. Notably, Gstue_Dup17 was not detected in Tanzania, indicating that this particular variant has not yet arrived in southern East Africa. However, further sampling points across the region would be required to confirm its presence and absence. Such studies could be facilitated by the association of tagging SNPs with these CNV variants, which provides an opportunity to design targeted assays, enabling higher resolution monitoring of their distribution and potential spread. In addition to the *Gste2* locus, we found there has been adaptive allele sharing at the *Cyp6aa/p* locus. Analysis revealed a single major CNV that is shared across all of the East African populations sampled. The widespread presence of this CNV highlights the potential of adaptive alleles to move across large geographical areas, overcoming barriers that otherwise restrict neutral gene flow. However, whether such barriers impact on the rate of spread would require a fine-scale longitudinal study targeting newly arisen or geographically restricted insecticide resistance variants. Potential targets include the gene duplications at the *Coejhe3e* locus, apparently under selection in northern Ethiopia, and the *Vgsc*-L995F allele, which is geographically restricted in East Africa and under selection in northernmost Ethiopia (Eukubay et al. 2026). Although CNV frequency was elevated at *Coejhe3e*, we observed no diplotype clusters with low heterozygosity, suggesting that these duplications have not yet become commonly associated with a single dominant diplotype background. Moreover, it has not yet been confirmed whether CNVs at the *Coejhe3e* locus impact insecticide resistance phenotypes in Ethiopia. Although *An. arabiensis* has historically been considered less impacted by insecticide resistance than *An. gambiae* (Kamau et al. 2008; Kawada et al. 2011; Protopopoff et al. 2013; Maliti et al. 2014; Nkya et al. 2014; Cisse et al. 2015; Mawejje et al. 2023; Ramaita et al. 2025), recent genomic and phenotypic evidence reveals that resistance alleles in *An. arabiensis* are geographically widespread and increasingly diverse across East Africa (Abdalla et al. 2014b; Yewhalaw et al. 2016; Alemayehu et al. 2017; Messenger et al. 2017; Hemming-Schroeder et al. 2018; Kiuru et al. 2018; Polo et al. 2025; Mutwiri et al. 2026). Our findings align with this emerging pattern, suggesting that resistance in *An. arabiensis* may be more extensive than previously recognised.

Our findings have important implications for the sustainability of malaria control in the region. The adaptive allele sharing we observed across East Africa implies that CNVs and SNPs conferring resistance to insecticides have a strong capacity to spread and reinvade areas upon elimination (The Anopheles gambiae 1000 Genomes Consortium 2017; Clarkson et al. 2021), even when populations are impacted by restricted gene flow. The transboundary movement of resistance alleles highlights that national vector control efforts to control the rise of insecticide resistance are vulnerable to re-invasion from neighbouring countries, as has been the case for vector elimination efforts in Mauritius and Sri Lanka (Aboobakar et al., 2012; WHO, 2023). Cross-country collaboration in insecticide resistance management is therefore essential. For example, initiatives include the Namibia-Angola Trans-Kunene Malaria Initiative (Khadka et al. 2018), the Southern Africa Malaria Elimination 8 programme (Sikaala et al. 2024) and recent Intergovernmental Authority on Development (IGAD) initiatives in East Africa (Mahmoud 2024). Such initiatives demonstrate the benefits of coordinated interventions, including harmonised policies, synchronised IRS campaigns and joint surveillance. Furthermore, high population connectivity across large geographical areas can provide a unique challenge for novel genetic control technologies such as gene drives, which face significant challenges from migration load (Noble et al. 2018; The Anopheles gambiae 1000 Genomes Consortium et al. 2020; Tajudeen et al. 2024). Nevertheless, the identification of isolated pockets such as those in the Central Ethiopian Rift Valley offers opportunities for their practical implementation, where the influx of wild-type alleles are less likely to dilute the frequency of the gene drive and the geographical spread of artificial genetic constructs could also be more easily controlled (North et al. 2013; Dhole et al. 2020). Furthermore, the identification of intermediate areas of gene flow, as we observed in southwestern Ethiopia, may provide areas for targeted vector control to prevent the spread of newly arisen or geographically restricted insecticide resistance variants. Prioritising these areas for routine monitoring with whole-genome sequencing would have the power to identify novel loci under selection and could therefore be used to provide an early warning of resistance alleles before they spread more broadly.

## Materials and Methods

### Mosquito sampling

This study used a total of 2662 *Anopheles arabiensis* whole genome sequences (WGS) available as part of the *Anopheles gambiae* 1000 Genome Project (https://malariagen.github.io/vector-data/ag3). This included 1,272 mosquitoes collected from fifteen sites across the northern, central, southern, southwestern, and western regions of Ethiopia, as previously described (Eukubay et al. 2026). Briefly, samples were collected between 2020 and 2023 from selected households using CDC light traps, Prokopack aspiration and manual aspiration. We also used WGS data from phase 3 (releases 3.0, 3.3, 3.7, 3.9 and 3.10) of the *Anopheles gambiae* 1000 Genome Project (Ag1000G) sampled in East Africa, including 512 from Kenya, 793 from Tanzania and 85 from Uganda (Figure 1). These samples were collected between 2006 and 2021 using methods previously described (The Anopheles gambiae 1000 Genomes Consortium 2021; Mwinyi et al. 2025; Polo et al. 2025).

### Whole-genome sequencing and data processing

The whole-genome sequencing of mosquitoes was performed using the Illumina HiSeq X platform as described previously (Eukubay et al. 2026). Bioinformatic analysis, including raw sequence processing, quality filtering, read alignment and variant calling, was conducted following the workflow developed by The Anopheles gambiae 1000 Genome Project (https://malariagen.github.io/vector-data/ag3/methods.html). Reads were mapped to the AgamP4 reference genome using BWA version 0.7.15, and single nucleotide polymorphism (SNP) data were produced with GATK version 3.7.0. Samples with a median coverage below 10x, those having less than 50% genome coverage, or exhibiting a high contamination level (>4.5%) were excluded. CNV calling was performed based on copy number states using normalised coverage data calculated in 300 bp genomic windows, and a Gaussian Hidden Markov Model (HMM) performed via hmmlearn (<u>GitHub - hmmlearn/hmmlearn:</u> <u>Hidden Markov Models in Python, with scikit-learn like API</u>) as previously detailed by (Lucas et al. 2019). The copy number of each gene was calculated as the median HMM value over all genomic windows covered by the gene. Samples were considered to have a CNV in these genes if the copy number was above the normal diploid copy number of 2. To enhance the accuracy of CNV prediction, individuals exhibiting high coverage variance (> 0.35) were excluded. CNV alleles were distinguished using reads aligning at the CNV breakpoints, as described in (Lucas et al. 2019). Gstue_Dup12 calls are part of the MalariaGEN data release. The other CNV allele was novel, and we identified diagnostic breakpoint reads *de-novo*. We named it Gstue_Dup17, and it was characterised by face-away read pairs aligning in the intervals 28596830-28597130 and 28605700-28606100, respectively, and individual reads soft-clipped at positions 28596832 or 28606222. This allele will be added to future releases of MalariaGEN.

Genotypes at biallelic SNPs that satisfied site filtering criteria were phased into haplotypes employing a combination of read-backed and statistical phasing techniques with WhatsHap V1.0 (Martin et al. 2016) and SHAPEIT V4.2 (Delaneau et al. 2019), respectively, adhering to the protocols of the *Anopheles gambiae* 1000 Genome Project https://malariagen.github.io/vector-data/ag3/methods.html).

### Population structure and genetic diversity

To investigate population connectivity, we constructed an unrooted neighbour-joining tree (NJT) and performed principal component analysis (PCA) across the 3L chromosome arm (15-41 Mbp), a region less affected by large structural rearrangements such as inversions and deletions (The Anopheles gambiae 1000 Genomes Consortium 2017). Analyses were based on 100,000 evenly distributed biallelic SNPs with a minimum allele count >2 and no missing calls. The NJT and PCA analyses were conducted using the malariagen_data python package https://malariagen.github.io/malariagen-data-python/latest/Ag3.html. We applied the elbow method using the <u>elbowplot</u> package (PyPI 2024) to the principal component scores (PC1 and PC2) to determine the optimal number of clusters, which was indicated *K=3* (Supplementary Figure S1). K-means clustering was then performed with the <u>scikit-learn</u> (Pedregosa et al., 2011) and <u>Matplotlib</u> Python packages (Hunter 2007). Averaged PC1 values were projected onto a geographic map using <u>GeoPandas</u> (Jordahl et al. 2020) and Plotly (Plotly Technologies Inc. n.d). To further support the PCA and NJT results, genetic differentiation was assessed using pairwise Hudson’s F_ST_ values (Hudson et al. 1992) between population cohorts organised by location and time of collection using the malariagen_data Python package. We used the F_ST_ estimates and a matrix of geographic distance calculated in the Python package Geopandas (Jordahl et al. 2020) to assess isolation by distance (IBD). To do this, we used a Mantel test (Mantel 1967) implemented in the <u>scikit-bio</u> Python package (Aton et al. 2025).

In addition to population connectivity analyses, we assessed genetic diversity both within Ethiopian cohorts and across East African *An. arabiensis*. Diversity analysis was restricted to the 3L chromosome arm (15-41 Mbp) (The Anopheles gambiae 1000 Genomes Consortium 2017), with a minimum cohort size of 10 individuals to ensure robust estimates. For Ethiopian samples, diversity summary statistics, including nucleotide diversity (𝛑), Watterson’s theta (𝛉w), and Tajima’s D, were calculated for each collection location and sampling season to evaluate temporal variation. A run of homozygosity (ROH) was performed to investigate the population history of outlier samples from the Southern Ethiopia (Dilla) site . Across East Africa, diversity analyses were performed by collection year, enabling comparisons of genetic variation over time and across geographic regions. All diversity statistics analyses were performed using the inbuilt Python packages from the malariagen_data <u>API</u>.

### Adaptive gene flow at insecticide-resistance loci

To investigate adaptive allele sharing across East Africa, we performed hierarchical clustering of diplotypes with city block genetic distance and complete linkage using the in-built function of the malariagen_data Python package. Diplotype clustering analyses were performed for insecticide resistance loci previously identified as under selection in Ethiopia (Eukubay et al. 2026), including the known metabolic resistance loci *Gste2* (AGAP009194) and the *Cyp6aa/p* gene cluster (AGAP002862-AGAP013128). We also investigated the *Coejhe1-5e* gene cluster (AGAP005833-AGAP005837), recently shown to be associated with organophosphate and pyrethroid insecticide resistance (Abdalla et al. 2014a; Omoke et al. 2024) and the target site locus *Vgsc* (AGAP004707) observed under selection in northern Ethiopia (Eukubay et al., 2026).

## Data availability

The sequences of the samples identified in this study were submitted to the European Nucleotide Archive (ENA; accession numbers are given in Supplementary Table S1).

## Acknowledgements

This study was supported by the MalariaGEN Vector Observatory which is an international collaboration working to build capacity for malaria vector genomic research and surveillance, and involves contributions by the following institutions and teams. Liverpool School of Tropical Medicine: Jon Brenas, Victoria Simpson, Lee Hart, Martin Donnelly; Wellcome Sanger Institute Alistair Miles, Eleanor Drury, Mara Lawniczak; The authors would like to thank the staff of the Wellcome Sanger Institute Sample Logistics, Sequencing, and Informatics facilities for their contributions.

## Author contributions

AE and KLB conducted the data analysis and interpretation and wrote the manuscript. HT and FG facilitated sample collection, processing, and data collection, with HT also supporting data interpretation. EL and AHK conducted data analysis. AM, DA, CCC and LG conceptualised and designed the study, interpreted the data and assisted in drafting the manuscript. All the authors revised the manuscript.

## References

Abbasi E. 2025. Climate Change and Vector-Borne Disease Transmission: The Role of Insect Behavioral and Physiological Adaptations. Integr Org Biol. 7(1):obaf011. 10.1093/iob/obaf011

Abdalla H et al. 2014a. Insecticide resistance in Anopheles arabiensis in Sudan: temporal trends and underlying mechanisms. Parasit Vectors. 7:213. 10.1186/1756-3305-7-213

Abdalla H et al. 2014b. Insecticide resistance in Anopheles arabiensis in Sudan: temporal trends and underlying mechanisms. Parasit Vectors. 7:213. 10.1186/1756-3305-7-213

Alemayehu E et al. 2017. Mapping insecticide resistance and characterization of resistance mechanisms in Anopheles arabiensis (Diptera: Culicidae) in Ethiopia. Parasit Vectors. 10(1):1–11. 10.1186/s13071-017-2342-y

Ashine T et al. 2024. Plasticity of blood feeding behavior of Anopheles mosquitoes in Ethiopia: a systematic review. Parasit Vectors. 17:408. 10.1186/s13071-024-06493-1

Ashine T et al. 2024. Anopheles arabiensis. Trends Parasitol. 40(1):91–92. 10.1016/j.pt.2023.08.011

Aton M et al. 2025. Scikit-bio: a fundamental Python library for biological omic data analysis. Nat Methods. [published online ahead of print]. 10.1038/s41592-025-02981-z

Barasa S et al. 2025. Assessing insecticide susceptibility status of Anopheles mosquitoes in Gondar zuria district, Northwest Ethiopia. Sci Rep. 15(1):14452. 10.1038/s41598-025-96370-3

Behrends GJ, Meheretu Y, Manthey JD. 2024. The Great Rift Valley is a more pronounced biogeographic barrier than the Blue Nile Valley for six Ethiopian Highland passerines in the eastern Afromontane biodiversity hotspot. Ornithology. 141(4). 10.1093/ornithology/ukae030

Bergey CM et al. 2020. Assessing connectivity despite high diversity in island populations of a malaria mosquito. Evol Appl. 13(2):417–431. 10.1111/eva.12878

Besansky NJ et al. 1997. Patterns of Mitochondrial Variation Within and Between African Malaria Vectors, Anopheles gambiae and An. arabiensis, Suggest Extensive Gene Flow. Genetics. 147(4):1817–1828. 10.1093/genetics/147.4.1817

Chanyalew T et al. 2022. Composition of mosquito fauna and insecticide resistance status of Anopheles gambiae sensu lato in Itang special district, Gambella, Southwestern Ethiopia. Malar J. 21(1):125. 10.1186/s12936-022-04150-5

Cisse MBM et al. 2015. Characterizing the insecticide resistance of Anopheles gambiae in Mali. Malar J. 14(1):327. 10.1186/s12936-015-0847-4

Clarkson CS et al. 2021. The genetic architecture of target-site resistance to pyrethroid insecticides in the African malaria vectors Anopheles gambiae and Anopheles coluzzii. Mol Ecol. 30(21):5303–5317. 10.1111/mec.15845

Delaneau O et al. 2019. Accurate, scalable and integrative haplotype estimation. Nat Commun. 10(1):24–29. 10.1038/s41467-019-13225-y

Denje T et al. 2023. A ground truthing analysis of climate, livelihood, and conflict in Turkana and West Pokot. [accessed 2026 Jan 31]. https://hdl.handle.net/10568/137994

Dhole S, Lloyd AL, Gould F. 2020. Gene Drive Dynamics in Natural Populations: The Importance of Density Dependence, Space, and Sex. Annu Rev Ecol Evol Syst. 51(1):505–531. 10.1146/annurev-ecolsys-031120-101013

Donnelly MJ, Licht MC, Lehmann T. 2001. Evidence for Recent Population Expansion in the Evolutionary History of the Malaria Vectors Anopheles arabiensis and Anopheles gambiae. Mol Biol Evol. 18(7):1353–1364. 10.1093/oxfordjournals.molbev.a003919

Eba K et al. 2021. Anopheles arabiensis hotspots along intermittent rivers drive malaria dynamics in semi-arid areas of Central Ethiopia. Malar J. 20(1):1–8. 10.1186/s12936-021-03697-z

Eligo N et al. 2024. Anopheles arabiensis continues to be the primary vector of Plasmodium falciparum after decades of malaria control in southwestern Ethiopia. Malar J. 23(1):14. 10.1186/s12936-024-04840-2

Enyew B, Hutjis R. 2015. Climate Change Impact and Adaptation in South Omo Zone, Ethiopia. J Geol Geophys. 4(3):1–14. 10.4172/2381-8719.1000208

Esayas E et al. 2024. Bionomic characterization of Anopheles mosquitoes in the Ethiopian highlands and lowlands. Parasit Vectors. 17(1):306. 10.1186/s13071-024-06378-3

Eukubay A et al. 2026. Spatial and temporal signatures of genomic insecticide resistance in the Anopheles arabiensis mosquito malaria vector from Ethiopia. Res Sq. rs.3.rs-8686648. 10.21203/rs.3.rs-8686648/v1

Freilich X et al. 2016. Comparative Phylogeography of Ethiopian anurans: impact of the Great Rift Valley and Pleistocene climate change. BMC Evol Biol. 16(1):206. 10.1186/s12862-016-0774-1

Gari T et al. 2016. Malaria incidence and entomological findings in an area targeted for a cluster-randomized controlled trial to prevent malaria in Ethiopia: results from a pilot study. Malar J. 15(1):145. 10.1186/s12936-016-1199-4

Getachew D, Balkew M, Tekie H. 2020. Anopheles larval species composition and characterization of breeding habitats in two localities in the Ghibe River Basin, southwestern Ethiopia. Malar J. 19(1):1–13. 10.1186/s12936-020-3145-8

Gimonneau G et al. 2012. Larval habitat segregation between the molecular forms of the mosquito Anopheles gambiae in a rice field area of Burkina Faso, West Africa. Med Vet Entomol. 26(1):9–17. 10.1111/j.1365-2915.2011.00957.x

Habtewold T et al. 2025. Gene-drive-capable mosquitoes suppress patient-derived malaria in Tanzania. Nature. 1–7. 10.1038/s41586-025-09685-6

Hemming-Schroeder E et al. 2018. Emerging Pyrethroid Resistance among Anopheles arabiensis in Kenya. Am J Trop Med Hyg. 98(3):704–709. 10.4269/ajtmh.17-0445

Hemming-Schroeder E et al. 2020. Ecological drivers of genetic connectivity for African malaria vectors Anopheles gambiae and An. arabiensis. Sci Rep. 10(1):19946. 10.1038/s41598-020-76248-2

Hudson RR, Slatkin M, Maddison WP. 1992. Estimation of levels of gene flow from DNA sequence data. Genetics. 132(2):583–589. 10.1093/genetics/132.2.583

Hunter JD. 2007. Matplotlib: A 2D Graphics Environment. Comput Sci Eng. 9(3):90–95. 10.1109/MCSE.2007.55

Jordahl K et al. 2020. geopandas/geopandas: v0.8.1. [accessed 2026 Jan 31]. https://zenodo.org/record/3946761. 10.5281/ZENODO.3946761

Kahamba NF et al. 2022. Using ecological observations to improve malaria control in areas where Anopheles funestus is the dominant vector. Malar J. 21:158. 10.1186/s12936-022-04198-3

Kamau L et al. 1999. Analysis of genetic variability in Anopheles arabiensis and Anopheles gambiae using microsatellite loci. Insect Mol Biol. 8(2):287–297. 10.1046/j.1365-2583.1999.820287.x

Kamau L et al. 2008. Status of insecticide susceptibility in Anopheles gambiae sensu lato and Anopheles funestus mosquitoes from Western Kenya. J Insect Sci. 8(1):11. 10.1673/031.008.1101

Kawada H et al. 2011. Distribution of a Knockdown Resistance Mutation (L1014S) in Anopheles gambiae s.s. and Anopheles arabiensis in Western and Southern Kenya. PLOS ONE. 6(9):e24323. 10.1371/journal.pone.0024323

Khadka A et al. 2018. Malaria control across borders: quasi-experimental evidence from the Trans-Kunene malaria initiative (TKMI). Malar J. 17(1):224. 10.1186/s12936-018-2368-4

Killeen GF, Govella NJ, Lwetoijera DW, Okumu FO. 2016. Most outdoor malaria transmission by behaviourally-resistant Anopheles arabiensis is mediated by mosquitoes that have previously been inside houses. Malar J. 15(1):225. 10.1186/s12936-016-1280-z

Kirby MJ, Lindsay SW. 2004. Responses of adult mosquitoes of two sibling species, Anopheles arabiensis and A. gambiae s.s. (Diptera: Culicidae), to high temperatures. Bull Entomol Res. 94(5):441–448. 10.1079/ber2004316

Kitau J et al. 2012. Species Shifts in the Anopheles gambiae Complex: Do LLINs Successfully Control Anopheles arabiensis? PLOS ONE. 7(3):e31481. 10.1371/journal.pone.0031481

Kiuru CW et al. 2018. Status of insecticide resistance in malaria vectors in Kwale County, Coastal Kenya. Malar J. 17:3. 10.1186/s12936-017-2156-6

Kreppel KS et al. 2020. Emergence of behavioural avoidance strategies of malaria vectors in areas of high LLIN coverage in Tanzania. Sci Rep. 10(1):14527. 10.1038/s41598-020-71187-4

Kweyamba PA et al. 2025. Contrasting vector competence of three main East African Anopheles malaria vector mosquitoes for Plasmodium falciparum. Sci Rep. 15(1):2286. 10.1038/s41598-025-86409-w

Lawrence TJ et al. 2023. Shifting climate zones and expanding tropical and arid climate regions across Kenya (1980–2020). Reg Environ Change. 23(2):59. 10.1007/s10113-023-02055-w

Lehmann T et al. 1996. Genetic differentiation of Anopheles gambiae populations from East and West Africa: comparison of microsatellite and allozyme loci. Heredity. 77(2):192–200. 10.1038/hdy.1996.124

Lucas ER et al. 2019. Whole-genome sequencing reveals high complexity of copy number variation at insecticide resistance loci in malaria mosquitoes. Genome Res. 29(8):1250–1261. 10.1101/gr.245795.118

Lucas ER et al. 2024. Copy number variants underlie the major selective sweeps in insecticide resistance genes in Anopheles arabiensis from Tanzania. PLoS Biol. 22(12):e3002898. 10.1371/journal.pbio.3002898

Lucas RE et al. 2023. Genome-wide association studies reveal novel loci associated with pyrethroid and organophosphate resistance in Anopheles gambiae and Anopheles coluzzii. Nat Commun. 14:4946. 10.1038/s41467-023-40693-0

Luhunga PM, Songoro AE. 2020. Analysis of Climate Change and Extreme Climatic Events in the Lake Victoria Region of Tanzania. Front Clim. 2. 10.3389/fclim.2020.559584

Lwetoijera DW et al. 2014. Increasing role of Anopheles funestus and Anopheles arabiensis in malaria transmission in the Kilombero Valley, Tanzania. Malar J. 13(1):331. 10.1186/1475-2875-13-331

Mahmoud Y. 2024. IGAD; [accessed 2026 Feb 4]. https://igad.int/progress-towards-malaria-elimination-in-the-igad-region-a-collaborative-approach/

Maliti D et al. 2014. Islands and Stepping-Stones: Comparative Population Structure of Anopheles gambiae sensu stricto and Anopheles arabiensis in Tanzania and Implications for the Spread of Insecticide Resistance. PLOS ONE. 9(10):e110910. 10.1371/journal.pone.0110910

Mantel N. 1967. The detection of disease clustering and a generalized regression approach. Cancer Res. 27(2):209–220

Martin M et al. 2016. WhatsHap : fast and accurate read-based phasing. 1–18. 10.1101/085050

Martinez-Torres D et al. 1998. Molecular characterization of pyrethroid knockdown resistance (kdr) in the major malaria vector Anopheles gambiae s.s. Insect Mol Biol. 7(2):179–184. 10.1046/j.1365-2583.1998.72062.x

Mawejje HD et al. 2023. Characterizing pyrethroid resistance and mechanisms in *Anopheles gambiae* (*s.s.*) and *Anopheles arabiensis* from 11 districts in Uganda. Curr Res Parasitol Vector-Borne Dis. 3:100106. 10.1016/j.crpvbd.2022.100106

Messenger LA et al. 2017. Insecticide resistance in Anopheles arabiensis from Ethiopia (2012–2016): a nationwide study for insecticide resistance monitoring. Malar J. 16:469. 10.1186/s12936-017-2115-2

Mitchell SN et al. 2014. Metabolic and target-site mechanisms combine to confer strong DDT resistance in Anopheles gambiae. PLoS ONE. 9(3). 10.1371/journal.pone.0092662

Msugupakulya BJ et al. 2023. Changes in contributions of different Anopheles vector species to malaria transmission in east and southern Africa from 2000 to 2022. Parasit Vectors. 16(1):1–16. 10.1186/s13071-023-06019-1

Müller P et al. 2008. Field-Caught Permethrin-Resistant Anopheles gambiae Overexpress CYP6P3, a P450 That Metabolises Pyrethroids. PLOS Genet. 4(11):e1000286. 10.1371/journal.pgen.1000286

Mustafa MSEK, Jaal Z, Abu Kashawa S, Mohd Nor SA. 2021. Population genetics of Anopheles arabiensis, the primary malaria vector in the Republic of Sudan. Malar J. 20(1):1–13. 10.1186/s12936-021-03994-7

Muturi EJ et al. 2010. Population Genetic Structure of Anopheles Arabiensis (Diptera: Culicidae) in a Rice Growing Area of Central Kenya. J Med Entomol. 47(2):144–151. 10.1093/jmedent/47.2.144

Mutwiri WK et al. 2026. Insecticide resistance profiles of Anopheles arabiensis and relationship with Microsporidia MB infection in two rice agroecosystems in Kenya. Parasit Vectors. 19(1):84. 10.1186/s13071-025-07212-0

Mwima R et al. 2023. Potential persistence mechanisms of the major Anopheles gambiae species complex malaria vectors in sub-Saharan Africa: a narrative review. Malar J. 22(1):336. 10.1186/s12936-023-04775-0

Mwima R et al. 2025. Assessing the population genetic structure and demographic history of Anopheles gambiae and An. arabiensis at island and mainland sites in Uganda: Implications for testing novel malaria vector control approaches. 2025.05.19.654785. 10.1101/2025.05.19.654785

Mwinyi SH. et al. 2025. Genomic Analysis Reveals a New Cryptic Taxon Within the Anopheles gambiae Complex With a Distinct Insecticide Resistance Profile in the Coast of East Africa. Mol Ecol. 34(10):e17762. 10.1111/mec.17762

Nagi SC et al. 2024. Parallel Evolution in Mosquito Vectors-A Duplicated Esterase Locus is Associated With Resistance to Pirimiphos-methyl in Anopheles gambiae. Mol Biol Evol. 41(7):1–11. 10.1093/molbev/msae140

Naidoo K, Oliver SV. 2025. Gene drives: an alternative approach to malaria control? Gene Ther. 32(1):25–37. 10.1038/s41434-024-00468-8

Ng’habi KR et al. 2011. Population genetic structure of Anopheles arabiensis and Anopheles gambiae in a malaria endemic region of southern Tanzania. Malar J. 10(1):289. 10.1186/1475-2875-10-289

Nkya TE et al. 2014. Insecticide resistance mechanisms associated with different environments in the malaria vector Anopheles gambiae: a case study in Tanzania. Malar J. 13(1):28. 10.1186/1475-2875-13-28

Noble C et al. 2018. Current CRISPR gene drive systems are likely to be highly invasive in wild populations. eLife. 7:e33423. 10.7554/eLife.33423

North A, Burt A, Godfray HCJ. 2013. Modelling the spatial spread of a homing endonuclease gene in a mosquito population. J Appl Ecol. 50(5):1216–1225. 10.1111/1365-2664.12133

Nyanjom SRG et al. 2003. Population Genetic Structure of Anopheles arabiensis Mosquitoes in Ethiopia and Eritrea. J Hered. 94(6):457–463. 10.1093/jhered/esg100

Odero JO et al. 2025. Genomic evidence of spatially structured gene flow and divergent insecticide resistance backgrounds of the malaria vector Anopheles funestus in Tanzania. Genetics. 230(4):iyaf117. 10.1093/genetics/iyaf117

Omoke D et al. 2024. Whole transcriptomic analysis reveals overexpression of salivary gland and cuticular proteins genes in insecticide-resistant Anopheles arabiensis from Western Kenya. BMC Genomics. 25(1). 10.1186/s12864-024-10182-9

Orondo PW et al. 2021. Insecticide resistance status of Anopheles arabiensis in irrigated and non-irrigated areas in western Kenya. Parasit Vectors. 14:335. 10.1186/s13071-021-04833-z

Pabst R et al. 2025. Global invasion patterns and dynamics of disease vector mosquitoes. Nat Commun. 16(1):9127. 10.1038/s41467-025-64446-3

Pedregosa F et al. 2011. Scikit-learn: Machine Learning in Python. Mach Learn Python. 12:2825–2830

Plotly Technologies Inc. n.d. Plotly; [accessed 2026 Feb 20]. https://plotly.com/python/

Polo B et al. 2025. Genomic surveillance reveals geographical heterogeneity and differences in known and novel insecticide resistance mechanisms in Anopheles arabiensis across Kenya. BMC Genomics. 26:599

Protopopoff N et al. 2013. High level of resistance in the mosquito Anopheles gambiae to pyrethroid insecticides and reduced susceptibility to bendiocarb in north-western Tanzania. Malar J. 12:149. 10.1186/1475-2875-12-149

PyPI. 2024. elbowplot: A simple library to plot the elbow plot for K-means clustering. [accessed 2026 Jan 31]. https://github.com/yourusername/elbowplot

Ramaita E et al. 2025. Insecticide resistance intensity in *Anopheles gambiae* (*s.l*.) from five malaria epidemiological zones in Kenya. Curr Res Parasitol Vector-Borne Dis. 7:100252. 10.1016/j.crpvbd.2025.100252

Riveron JM et al. 2014. A single mutation in the GSTe2 gene allows tracking of metabolically based insecticide resistance in a major malaria vector. Genome Biol. 15(2):R27. 10.1186/gb-2014-15-2-r27

Sikaala CH et al. 2024. Malaria elimination and the need for intensive inter-country cooperation: a critical evaluation of regional technical co-operation in Southern Africa. Malar J. 23:62. 10.1186/s12936-024-04891-5

Strauss AT et al. 2020. Vector demography, dispersal and the spread of disease: Experimental epidemics under elevated resource supply. Funct Ecol. 34(12):2560–2570. 10.1111/1365-2435.13672

Tajudeen YA et al. 2024. Transforming malaria prevention and control: the prospects and challenges of gene drive technology for mosquito management. Ann Med. 55(2):2302504. 10.1080/07853890.2024.2302504

Tarekegn M, Tekie H, Wolde-hawariat Y, Dugassa S. 2022. Habitat characteristics and spatial distribution of Anopheles mosquito larvae in malaria elimination settings in Dembiya District, Northwestern Ethiopia. Int J Trop Insect Sci. 42(4):2937–2947. 10.1007/s42690-022-00821-7

Temu E, Yan G. 2005. Microsatellite and mitochondrial genetic differentiation of Anopheles arabiensis (Diptera: Culicidae) from Western Kenya, the Great Rift Valley and coastal Kenya. Am J Trop Med Hyg. 73:726–33. 10.4269/ajtmh.2005.73.726

The Anopheles gambiae 1000 Genomes Consortium. 2017. Genetic diversity of the African malaria vector Anopheles gambiae. Nature. 552(7683):96–100. 10.1038/nature24995

The Anopheles gambiae 1000 Genomes Consortium et al. 2020. Genome variation and population structure among 1142 mosquitoes of the African malaria vector species Anopheles gambiae and Anopheles coluzzii. Genome Res. 30(10):1533–1546. 10.1101/gr.262790.120

The Anopheles gambiae 1000 Genomes Consortium. 2021. Genome variation and population structure in three African malaria vector species within the *Anopheles gambiae* complex. Manubot. [accessed 2026 Feb 1]. https://malariagen.github.io/ag1000g-phase3-data-paper/

Wiebe A et al. 2017. Geographical distributions of African malaria vector sibling species and evidence for insecticide resistance. Malar J. 16(1):85. 10.1186/s12936-017-1734-y

Worku MA, Feyisa GL, Beketie KT. 2022. Climate trend analysis for a semi-arid Borana zone in southern Ethiopia during 1981–2018. Environ Syst Res. 11(1):2. 10.1186/s40068-022-00247-7

Woyessa D, Yewhalaw D. 2025. Anopheles mosquito fauna, blood meal sources and transmission intensity from high and moderate malaria endemic areas of Ethiopia. Sci Rep. 15(1):10636. 10.1038/s41598-025-94739-y

Yewhalaw D, Kweka EJ, Yewhalaw D, Kweka EJ. 2016. Insecticide Resistance in East Africa — History, Distribution and Drawbacks on Malaria Vectors and Disease Control. In: Insecticides Resistance. IntechOpen [accessed 2026 Jan 29]. https://www.intechopen.com/chapters/49270. 10.5772/61570

Zhou G et al. 2022. Irrigation-Induced Environmental Changes Sustain Malaria Transmission and Compromise Intervention Effectiveness. J Infect Dis. 226(9):1657–1666. 10.1093/infdis/jiac361

