## Supplemental Tables and Figures for "Genomic connectivity and the spread of adaptive insecticide resistance alleles in *Anopheles arabiensis* from East Africa"

**Supplementary Files**

**Supplementary Table S1**. List of sample sets and accession numbers of the sequence samples. Samples with multiple run accessions are given within their respective sample sets.

| **Sample sets** | **Accession number** |
| --- | --- |
| 1270-VO-MULTI-PAMGEN-VMF00162 | ERR6045394-ERR6045422, ERR6045425-ERR6045453,ERR6045456-ERR6045484 |
| 1270-VO-MULTI-PAMGEN-VMF00162 | ERR5987917-ERR5987948,ERR5987949-ERR5987980,ERR5987981-ERR5988012 |
| 1270-VO-MULTI-PAMGEN-VMF00162 | ERR5967771-ERR5967798,ERR5967799-ERR5967826,ERR5967827-ERR5967854 |
| 1270-VO-MULTI-PAMGEN-VMF00218 | ERR10419623-ERR10419646 |
| 1270-VO-MULTI-PAMGEN-VMF00218 | ERR10419672-ERR10419694 |
| 1270-VO-MULTI-PAMGEN-VMF00218 | ERR10490762-ERR10490785 |
| 1270-VO-MULTI-PAMGEN-VMF00218 | ERR10490810-ERR10490977 |
| 1270-VO-MULTI-PAMGEN-VMF00218 | ERR10970454-ERR10970494,  ERR11041589-ERR11041629 |
| 1324-VO-ET-GOLASSA-VMF00257 | ERR12262329-ERR12262396 |
| 1324-VO-ET-GOLASSA-VMF00257 | ERR12314213-ERR12314278 |
| 1324-VO-ET-GOLASSA-VMF00257 | ERR12325996-ERR12326278 |
| 1324-VO-ET-GOLASSA-VMF00257 | ERR12547479-ERR12547495 and ERR12767674 |
| 1324-VO-ET-GOLASSA-VMF00275 | ERR12871282-ERR12871307 |
| 1324-VO-ET-GOLASSA-VMF00275 | ERR12948031-ERR12948246 |
| 1324-VO-ET-GOLASSA-VMF00275 | ERR12983126-ERR12983194 |
| 1324-VO-ET-GOLASSA-VMF00275 | ERR13101382-ERR13101590 |
| 1324-VO-ET-GOLASSA-VMF00275 | ERR13146891-ERR13146976 |

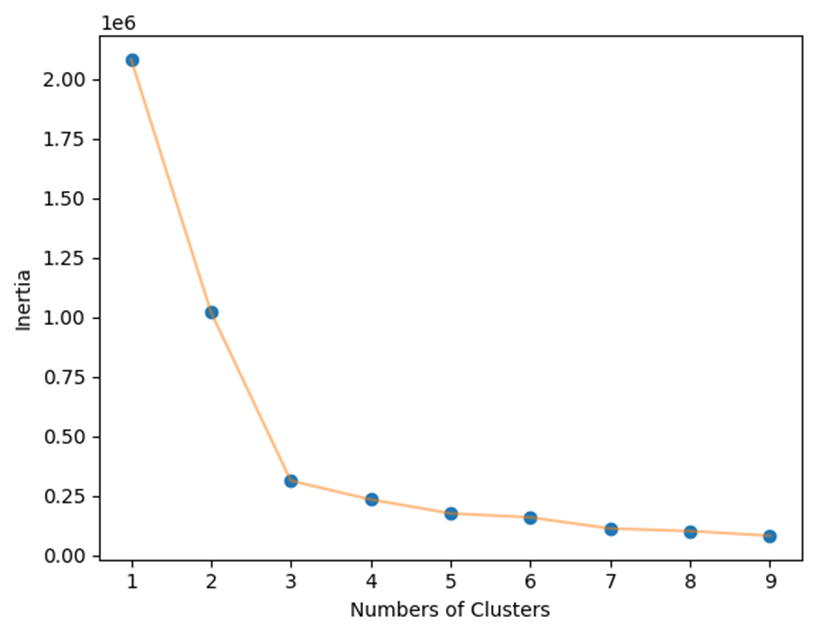

**Supplementary Figure S1.** Elbow method plot showing the relationship between the number of clusters (K) and the within-cluster sum of squares (inertia) for the PCA dataset. The sharp decrease at K=3 followed by a gradual decline, indicates three clusters as the optimal number for subsequent KMeans clustering.

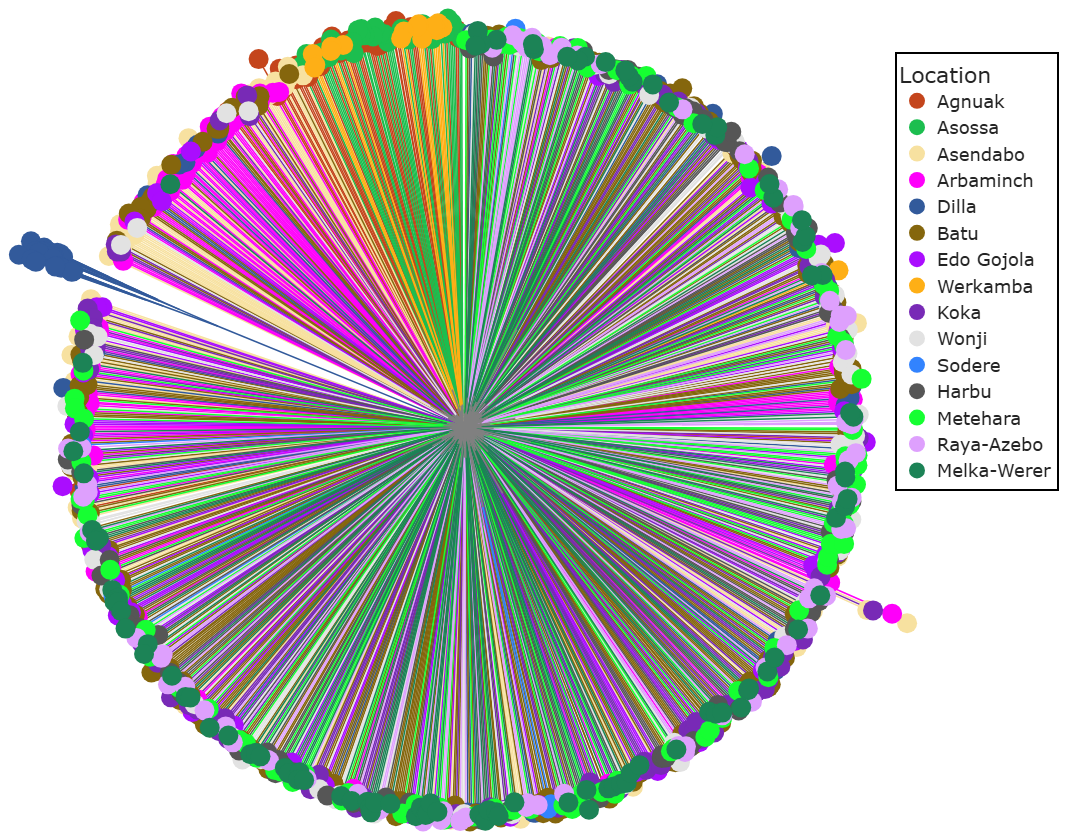

**Supplementary S2**. Neighbour-joining tree of *Anopheles arabiensis* coloured by location in Ethiopia using 100,000 SNPs on chromosome 3L.

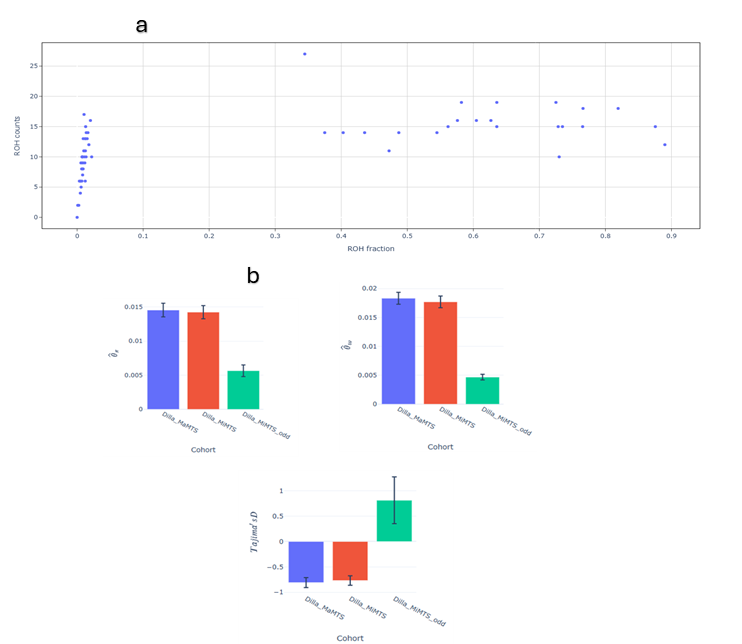

**Supplementary S3.** (a) Runs of Homozygosity (ROH) plot and (b) a bar plot of nucleotide diversity, Watterson’s estimator, and Tajima’s D comparing individuals of *An. arabiensis* from Dilla collected during the Major Malaria Transmission Season (MaMTS), Minor Malaria Transmission Season (MiMTS) and for individuals which were outlier samples on the PCA and NJT.

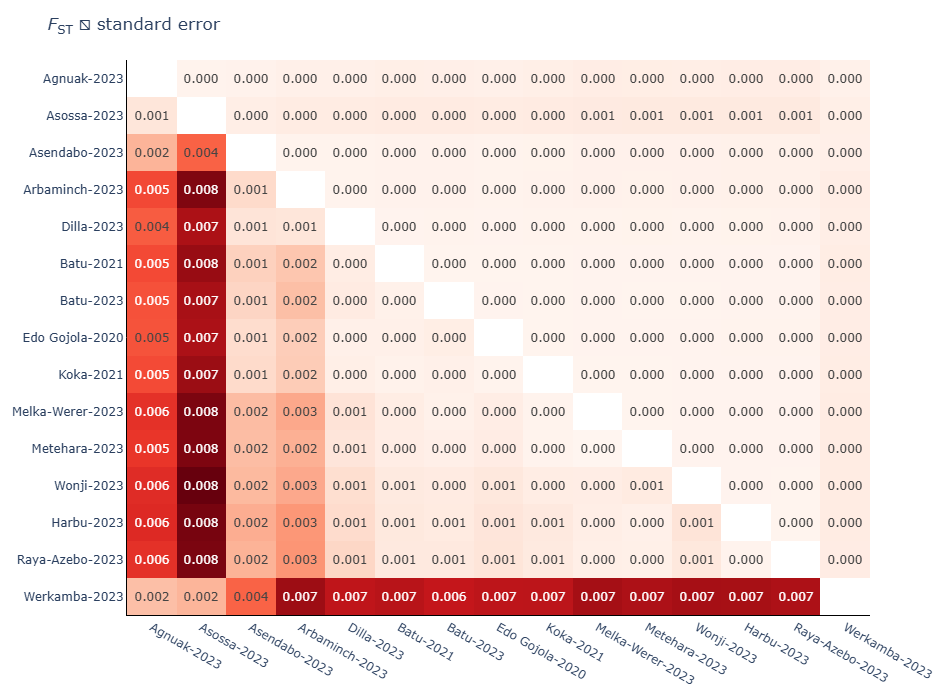

**Supplementary S4**. Measures of pairwise F_ST_ of *Anopheles arabiensis* from different sampling locations and years. The lower triangle matrix shows pairwise F_ST_ values, and the upper triangle shows standard error (SE).

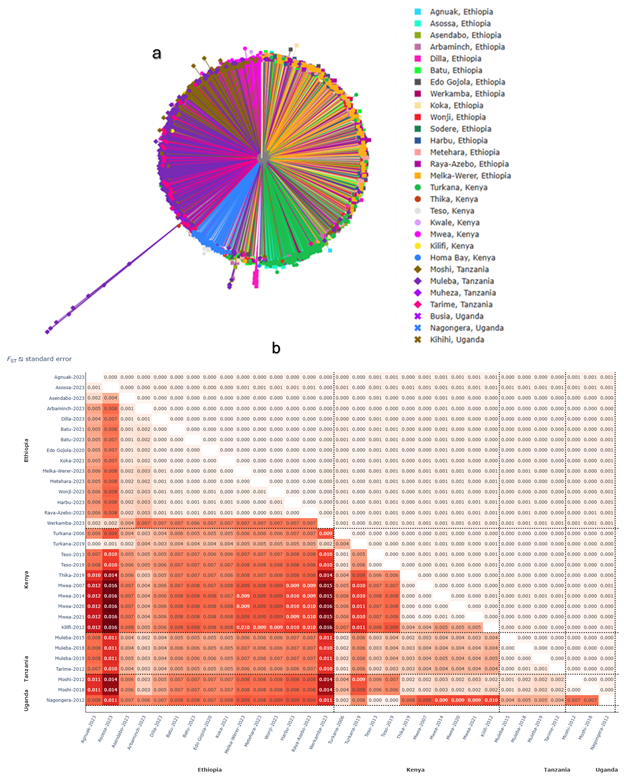

**Supplementary S5**. Population structure of Anopheles arabiensis across East Africa. (**a**) a neighbour-joining tree (NJT) plotted by location and country and (**b**) measures of fixation index (F_ST_) showing genetic differentiation of East African *Anopheles arabiensis* by location and year. The lower triangle displays pairwise FST, and the upper triangle shows standard error (SE).

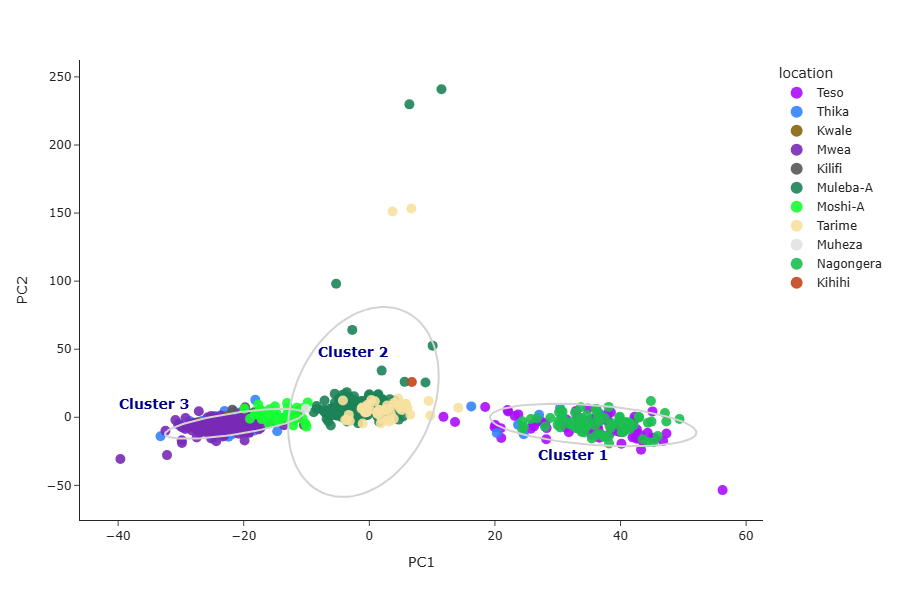

**Supplementary Figure S6.** Principal component analysis (PCA) of *Anopheles arabiensis* populations from East Africa (Kenya excluding Turkana, Tanzania, and Uganda). Each point represents an individual genome, colour-coded by sampling location. The three clusters are determined by K-means clustering, with ellipses added to illustrate the spread of points in each cluster and scaled to represent the 95% confidence interval.

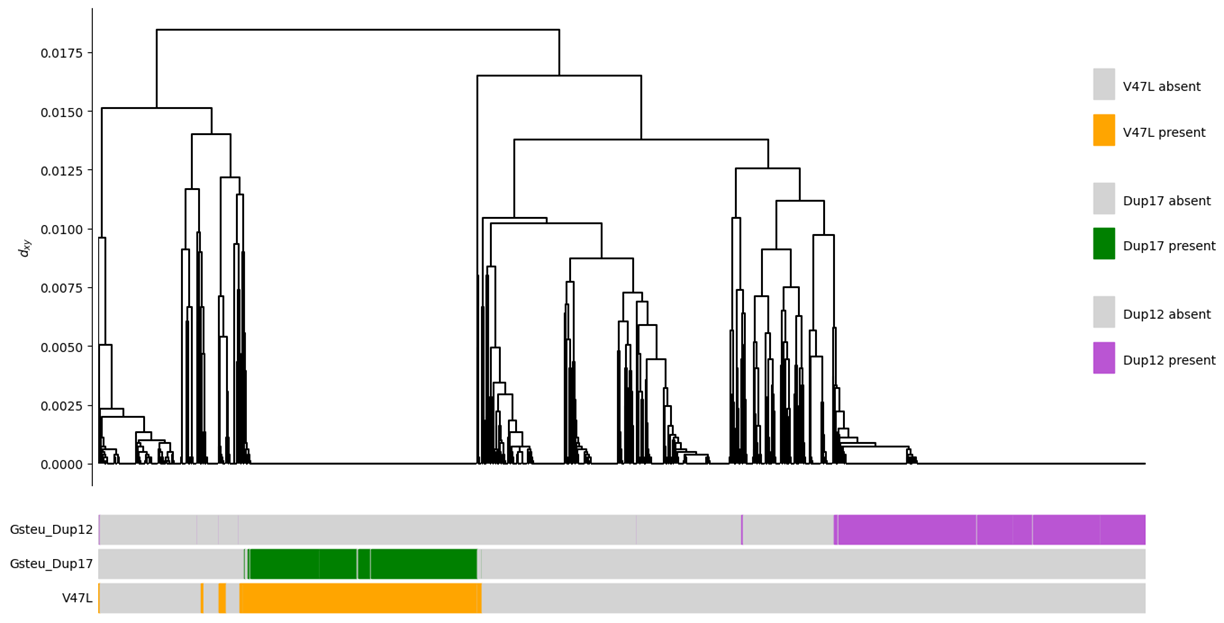

**Supplementary Figure S7**. Haplotype clustering of the genomic region surrounding *Gste2* in Ethiopian *Anopheles arabiensis*. Haplotypes carrying the CNV duplications Gstue-Dup17 with its associated SNP (V47L) and Gstue-Dup12 are annotated under the dendrogram as colour bars indicating presence and absence.

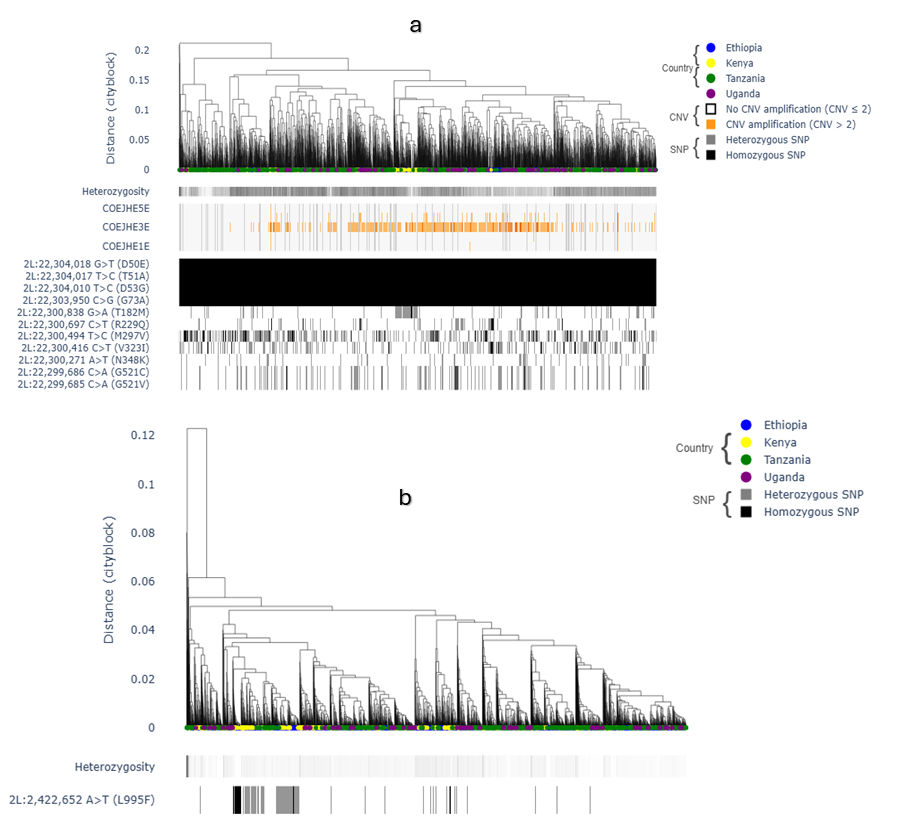

**Supplementary Figure S8**. Diplotype clustering of East African *Anopheles arabiensis* at (**a**) the *Coejhe1-5e* and (**b**) *Vgsc* genes. Heterozygosity and copy number variation (CNV) are shown as colour bars below the dendrogram, with darker shades on the heterozygosity bar representing higher diversity and orange-red color gradients on the copy number bar showing CNV amplification. Non-synonymous SNPs are plotted below the copy number bar, with a gray-to-black color gradient, showing heterozygous and homozygous non-synonymous SNPs, respectively.
